# Clonal hematopoiesis-associated DNMT3A mutations derepress bivalent Polycomb target genes and enhance macrophage migration

**DOI:** 10.64898/2026.09.24.754138

**Authors:** Yuri Lee, Philip K. Farahat, Jennifer P. Nguyen, Yoshiko Takahashi, Jung-Yeon Lim, Xin Huang, Camille Brzechffa, Marie E. S. Wheeler, Hye-Ju Yang, Jihye Lee, Zachary S. Pope, Daniella J. Lu, Akshata N. Rudrapatna, Alec P. Pankow, Xiaoting Chen, Hongseok Yun, Matthew T. Weirauch, Ivan Marazzi, Brad R. Rosenberg, Emily R. Miraldi, Eirini P. Papapetrou, S. Armando Villalta, Angela G. Fleischman, Minji Byun

## Abstract

Somatic mutations in the de novo DNA methyltransferase *DNMT3A* are frequent in clonal hematopoiesis, but how they alter the function of differentiated myeloid cells remains unclear. Here, we used isogenic human embryonic stem cell-derived macrophage models to define the consequences of DNMT3A dysfunction during myeloid differentiation. *DNMT3A*-mutant macrophages selectively upregulated bivalent Polycomb target genes, accompanied by DNA hypomethylation, reduced H3K27me3 and increased promoter accessibility. Despite broad epigenetic changes across bivalent promoters, transcriptional derepression occurred at a subset of loci characterized by lower baseline DNA methylation and higher H3K27me3, identifying pre-existing chromatin state as a determinant of transcriptional response. This Polycomb-associated program was conserved in murine macrophages and enriched for genes involved in cell migration and wound repair. *DNMT3A*-mutant macrophages exhibited enhanced migration and preferential early recruitment to injured tissue. These findings link clonal hematopoiesis-associated epigenetic alterations to selective transcriptional reprogramming and altered function of differentiated myeloid cells.

## Introduction

Somatic mutations in epigenetic regulators are common in hematologic malignancies and age-associated hematopoietic clonal expansions, but how they produce lasting changes in gene regulation and function in differentiated immune cells remains unclear. The *de novo* DNA methyltransferase DNMT3A plays a central role in establishing DNA methylation patterns during development and hematopoiesis.^1,2^ Recurrent *DNMT3A* mutations occur in acute myeloid leukemia and related myeloid neoplasms and are the most frequent lesions in clonal hematopoiesis of indeterminate potential (CHIP), in which mutated hematopoietic stem and progenitor cells clonally expand with age.^3–6^ Individuals with *DNMT3A*-mutant clonal hematopoiesis have an increased risk of hematologic cancers and non-malignant age-related disorders affecting various organs, highlighting the systemic impact of perturbing *de novo* DNA methylation in the hematopoietic system.^4,5,7^

Myeloid cells, particularly monocytes and macrophages, are key effectors of tissue homeostasis and inflammation and have emerged as critical mediators of CHIP-associated pathology.^7–12^ Transcriptomic analyses of *DNMT3A*-mutant myeloid cells have revealed widespread changes in gene expression, including altered inflammatory and stress-response programs.^12–18^ In parallel, DNA methylation profiling of *DNMT3A*-mutant hematopoietic cells has identified characteristic hypomethylation at large domains and regulatory elements.^13,18–23^ However, directly linking these epigenetic alterations to specific transcriptional outputs has proven challenging. An outstanding question is how *DNMT3A* mutations acquired in stem and progenitor cells are translated into stable, locus-specific changes in chromatin and gene regulation in their differentiated myeloid progeny, and whether such changes are sufficient to alter myeloid cell behavior. Addressing this requires systems that isolate the cell-intrinsic effects of *DNMT3A* mutation from inflammatory or environmental cues, and that allow systematic mapping of chromatin state and gene expression in mutant versus wild-type myeloid cells.

Here, we used isogenic human embryonic stem cells (hESCs) engineered to carry *DNMT3A* mutations and differentiated them into macrophages to define the cell-intrinsic consequences of DNMT3A disruption on chromatin landscapes, transcriptional programs, and macrophage behavior. We integrated these data with complementary murine models in which *Dnmt3a* is mutated in hematopoietic stem and progenitor cells and *in vivo* assays of myeloid cell recruitment to tissue injury. This approach allowed us to identify promoter chromatin features that predispose genes to dysregulation in *DNMT3A*-mutant macrophages and to link these molecular changes to altered migratory and injury-response behaviors *in vivo*, thereby connecting somatic epigenetic lesions in stem cells to durable functional reprogramming of innate immune cells.

## Results

### Polycomb Repressive Complex 2 target genes are upregulated in macrophages derived from *DNMT3A*-mutant hESCs

Most *DNMT3A* mutations associated with clonal hematopoiesis are heterozygous loss-of-function or dominant-negative alleles, such as the R882H mutation.^24–26^ To model clonal hematopoiesis-relevant DNMT3A defects, we engineered hESC clones harboring either a monoallelic truncation mutation (HET: *DNMT3A*^+/-^) or a monoallelic dominant-negative mutation (R882H: *DNMT3A*^+/R882H^). hESC clones subjected to the same gene editing procedures but lacking mutations at the targeted loci served as wild-type controls (WT: *DNMT3A*^+/+^)^27^ (**Fig. 1a**). Macrophages were derived from these clonal hESCs under defined *in vitro* conditions using an established differentiation protocol.^13^

**Fig. 1.**
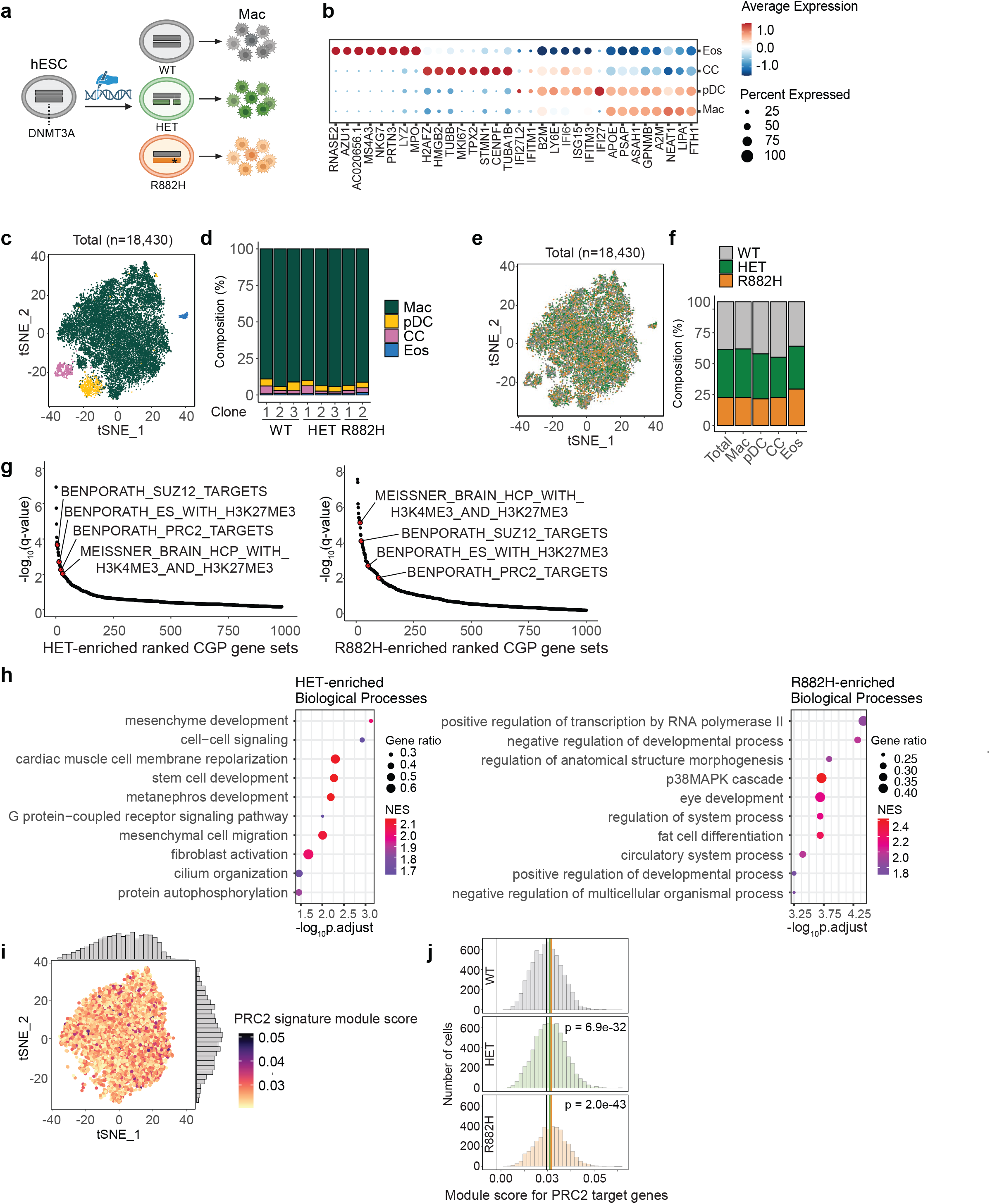
*DNMT3A* mutations drive derepression of PRC2 target genes in hESC-derived macrophages. **a,** Schematic of the study design using hESC-derived macrophages. A hand holding a pen denotes CRISPR-Cas9-mediated gene editing. Mac, macrophage. Created with BioRender.com. **b**, Dot plot showing expression of canonical marker genes across clusters. Cluster identities are shown on the y-axis and marker genes on the x-axis. Circle size indicates the percentage of cells in each cluster expressing the gene; color intensity indicates the mean scaled expression among expressing cells. Mac, macrophages; pDC, plasmacytoid dendritic cell (pDC)-like cells; CC, cycling cells; Eos, eosinophil-like cells. **c,** t-SNE projection of scRNA-seq data from hESC-derived macrophages, including three WT clones, three *DNMT3A-*HET (HET) clones, and two *DNMT3A*-R882H (R882H) clones. The total number of cells analyzed is indicated. **d,** Proportion of cells of each genotype assigned to each cluster. **e,** t-SNE plot colored by genotype (WT, HET, and R882H) showing the contribution of each genotype to all four clusters. **f**, Proportion of each cluster accounted for by the three genotypes. **g,** Gene set enrichment analysis (GSEA) of Chemical and Genetic Perturbation (CGP) gene sets from MSigDB.^28^ PRC2-related pathways (highlighted in red) rank among the most significantly enriched. Each dot represents a gene set ranked by adjusted p-value (Padj) of enrichment. Top 1,000 ranked pathways are shown for clarity. **h**, GSEA of Gene Ontology Biological Process (GO:BP) gene sets from MSigDB. Redundant, semantically similar terms were collapsed using the simplify() function in clusterProfiler^58^ (semantic similarity measured by the Wang method, similarity cutoff = 0.65). The top 10 pathways, ranked by Padj, are shown. **i**, t-SNE plots of the top 50% scoring cells in the macrophage cluster (WT, HET, R882H, and all genotypes combined), colored by PRC2 module score (heat map) calculated with UCell^60^ using a custom PRC2 target gene set (Supplementary Table 7, Cluster_1) defined by high promoter H3K27me3 in hESC-derived macrophages. **j.** Distribution of PRC2 target gene module scores in WT, HET, and R882H cells. Statistical significance, calculated by a Wilcoxon rank-sum test, is shown.

Single-cell RNA sequencing (scRNA-seq) at day 33 of differentiation revealed a dominant macrophage population and three minor subsets corresponding to plasmacytoid dendritic cell (pDC)-like cells, cycling cells, and eosinophil-like cells (**Fig. 1b–d**, and **Supplementary Table 1**). Macrophages constituted 89-94% of total cells across all clones (**Fig. 1d** and **Supplementary Table 2**). All four clusters included cells from WT, HET, and R882H clones in proportions reflecting the overall cell distribution, and no cluster was clearly associated with a single genotype (**Fig. 1e,f** and **Supplementary Table 3**), indicating that clonal hematopoiesis-associated *DNMT3A* mutations do not generate genotype-specific cell populations under these conditions.

To gain insights into pathways altered in mutant macrophages, we performed Gene Set Enrichment Analysis (GSEA) using Chemical and Genetic Perturbation (CGP) gene sets from the Molecular Signatures Database (MSigDB).^28^ Gene sets representing targets of Polycomb Repressive Complex 2 (PRC2) core components (“…SUZ12_TARGETS” and “…PRC2_TARGETS”) and genes marked by PRC2-deposited H3K27me3 (“…H3K27ME3”) were among the most enriched in the mutant macrophages compared to WT (**Fig. 1g** and **Supplementary Table 4, Supplementary Table 5**). PRC2 targets are characteristically associated with developmental gene regulation.^29,30^ Consistent with this, GSEA of Gene Ontology (GO) Biological Processes revealed significant enrichment of development-related pathways in the mutant macrophages relative to WT macrophages (**Fig. 1h** and **Supplementary Table 6**).

To determine whether this PRC2 signature arose from a discrete subpopulation within the macrophage compartment, we calculated a module score based on a custom PRC2 target gene set defined by high promoter H3K27me3 levels in hESC-derived macrophages (**Supplementary Table 7**, Cluster_1). Cells with high module scores were evenly distributed throughout the macrophage population on the t-SNE map (**Fig. 1i**), indicating that the PRC2 target signature represents a general feature of macrophages rather than a restricted subpopulation.

Comparison of module score distributions revealed significantly elevated PRC2 module scores in both HET and R882H macrophages relative to WT (**Fig. 1j**), corroborating the GSEA findings.

### Bivalent chromatin state marks genes derepressed by *DNMT3A* mutations

We next asked whether genes differentially expressed in *DNMT3A*-mutant macrophages are associated with PRC2-linked chromatin states. PRC2 targets are typically lowly expressed and therefore prone to dropout in single-cell datasets. Moreover, *DNMT3A*-mutant and WT cultures showed highly similar cellular composition, dominated by macrophages (**Fig. 1d**), indicating that bulk measurements should predominantly capture macrophage-intrinsic transcriptional differences. We therefore performed high-depth bulk RNA-seq on isogenic *DNMT3A*-mutant and WT macrophages to improve detection of low-abundance transcripts and to link differential expression to reference chromatin-state annotations.

CGP GSEA of bulk RNA-seq recapitulated the results from scRNA-seq, demonstrating enrichment of PRC2-related gene sets in *DNMT3A*-mutant macrophages (**Extended Data Fig. 1a** and **Supplementary Table 8**). We then overlaid differentially expressed genes (DEGs) (**Supplementary Table 9**) from bulk RNA-seq onto monocyte chromatin-state maps from the Roadmap Epigenome Project.^31^ DEGs in mutant macrophages were most highly enriched for bivalent states (TssBiv, BivFlnk, EnhBiv) (**Fig. 2a** and **Supplementary Table 10**). Bivalent chromatin states are marked by both the activating modification H3K4me3 and the repressive mark H3K27me3, poising genes for either activation or silencing.^32^ DEGs were also enriched, to a lesser extent, for Polycomb-repressed chromatin (ReprPC), characterized by H3K27me3 in the absence of H3K4me3 (**Fig. 2a** and **Supplementary Table 10**). In contrast, conventional active promoter states (TssA), typically H3K4me3-positive and H3K27me3-negative, were depleted among DEGs (**Fig. 2a** and **Supplementary Table 10**). These data suggest that genes responsive to *DNMT3A* mutation are preferentially embedded in bivalent and Polycomb-repressed chromatin domains.

**Fig. 2.**
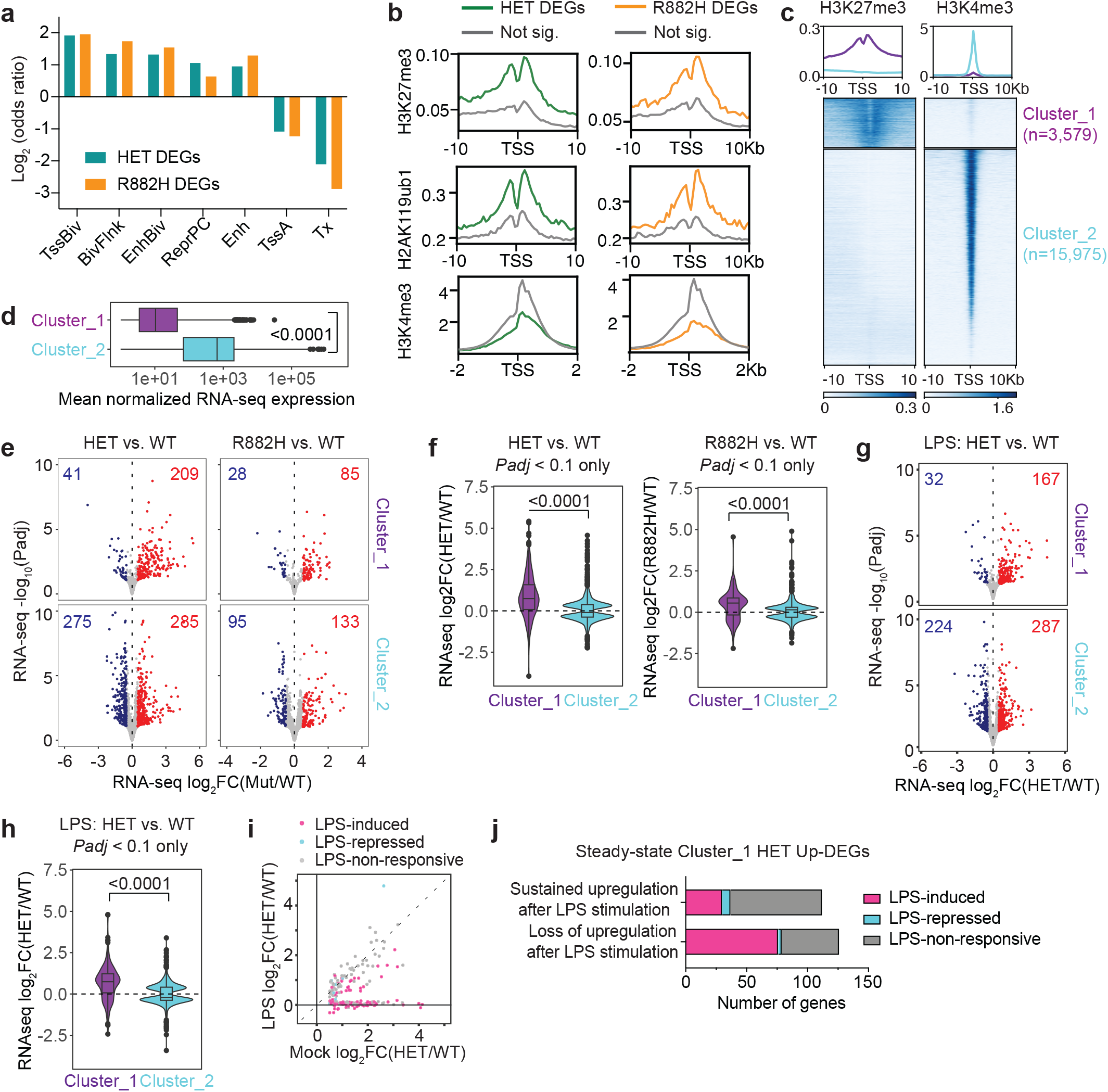
*DNMT3A* mutations preferentially increase expression of bivalent and Polycomb-repressed genes. **a,** Enrichment of monocyte chromatin states among DNMT3A-HET or DNMT3A-R882H differentially expressed genes (DEGs; Padj < 0.1 and |log2FC| > 0.5) compared with promoters of non-significant genes, assessed by two-sided Fisher’s exact test. TssBiv, bivalent/poised transcription start site (TSS); BivFlnk, flanking bivalent TSS/enhancer; EnhBiv, bivalent enhancer; ReprPC, Polycomb-repressed; Enh, enhancer; TssA, active TSS; Tx, strong transcription. **b,** Distribution of H3K27me3, H2AK119ub1, and H3K4me3 at TSSs and surrounding regions for HET or R882H DEGs, based on CUT&RUN data from WT hESC-derived macrophages. Not sig., non-significant genes. **c**, Heatmap of H3K27me3 and H3K4me3 signal within a 20-kb window centered on TSSs of expressed genes in WT hESC-derived macrophages, clustered by H3K27me3 signal to define H3K27me3-high bivalent promoters (Cluster_1) and H3K27me3-low promoters (Cluster_2) by k-means clustering (k = 2) with deepTools.^70^ The number of genes in each cluster is indicated. **d,** Comparison of mean normalized RNA-seq expression (DESeq2 baseMean)^63^ between Cluster_1 and Cluster_2 genes. Boxes show the median and interquartile range (IQR); whiskers extend to 1.5 × IQR. *P* value was calculated by Student’s *t*-test. **e,** Volcano plots showing shrunken log2FC (calculated using the lfcShrink function (DESeq2) with the apeglm shrinkage estimator) for HET versus WT and R882H versus WT under mock conditions, plotted against −log10(Padj). Each dot represents a gene; red dots indicate upregulated DEGs (Padj < 0.1 and log2FC > 0.5), whereas blue dots indicate downregulated DEGs (Padj < 0.1 and log2FC < −0.5). The numbers of upregulated and downregulated DEGs are indicated. **f,** Violin plots of shrunken log2FC (HET versus WT or R882H versus WT) for genes with Padj < 0.1 in Cluster_1 and Cluster_2 under mock conditions, illustrating the bias toward upregulation among Cluster_1 genes. *P* values were calculated by a Wilcoxon rank-sum test. **g,** Volcano plots of shrunken log2FC (HET versus WT) under LPS-stimulated conditions (100ng/mL, 4 h), plotted against -log10(Padj). Each dot represents a gene; red dots indicate upregulated DEGs (Padj < 0.1 and log2FC > 0.5), whereas blue dots indicate downregulated DEGs (Padj < 0.1 and log2FC < −0.5). The numbers of upregulated and downregulated DEGs are indicated. **h**, Violin plots of shrunken log2FC (HET versus WT) for genes with Padj < 0.1 in Cluster_1 and Cluster_2 under LPS-stimulated conditions, illustrating the bias toward upregulation among Cluster_1 genes. *P* values were calculated by a Wilcoxon rank-sum test. **i**, Relationship between HET versus WT shrunken log2FC under mock and LPS-stimulated conditions for Cluster_1 genes upregulated in HET macrophages under mock conditions (Padj < 0.1 and log2FC > 0.5). Dots are colored according to transcriptional response to LPS: LPS-induced, LPS versus mock in WT macrophages, Padj < 0.1 and log2FC > 1; LPS-repressed, Padj < 0.1 and log2FC < −1; LPS-non-responsive, all remaining genes. **j**, Cluster_1 genes upregulated in HET versus WT macrophages under mock conditions (Padj < 0.1 and log2FC > 0.5), grouped according to whether they remained upregulated or were no longer differentially expressed under LPS-stimulated conditions. Each group is further classified based on transcriptional response to LPS as LPS-induced, LPS-repressed, or LPS-non-responsive, as defined in **i**.

Because the Roadmap Epigenome annotations are derived from blood monocytes rather than the hESC-derived macrophages used here, we next asked whether a similar chromatin context was evident in our system. To this end, we used CUT&RUN to profile H3K27me3 and H3K4me3, as well as H2AK119ub1 deposited by Polycomb Repressive Complex 1 (PRC1), in macrophages derived from WT hESC clones. We then compared promoter enrichment of these marks between HET- and R882H-DEGs and genes not significantly altered in either mutant.

Promoters of DEGs exhibited higher levels of H3K27me3 and H2AK119ub1 and lower H3K4me3 compared to promoters of genes not significantly altered in either mutant (**Fig. 2b**), consistent with DNMT3A-responsive genes being enriched for promoters with bivalent and Polycomb-repressed features.

To systematically examine bivalent and Polycomb-repressed gene expression, we used the macrophage CUT&RUN data to cluster all expressed genes based on promoter H3K27me3 enrichment. This analysis identified two groups: H3K27me3-high genes (Cluster_1) and H3K27me3-low genes (Cluster_2) (**Fig. 2c, Supplementary Table 7**). H3K27me3-high genes displayed strongly reduced but detectable H3K4me3 at promoters (**Fig. 2c**) and were expressed at significantly lower levels than H3K27me3-low genes (**Fig. 2d**), a profile characteristic of bivalent promoters. We therefore refer to H3K27me3-high genes (Cluster_1) as bivalent Polycomb target genes in this system.

Strikingly, among Cluster_1 genes, most DEGs were upregulated in *DNMT3A*-mutant macrophages, whereas DEGs among H3K27me3-low (Cluster_2) genes showed a more balanced mix of up- and downregulation (**Fig. 2e,f**). We further investigated whether this preferential derepression of bivalent genes persists upon macrophage activation, using lipopolysaccharide (LPS)-stimulated RNA-seq data generated in WT and HET macrophages (R882H LPS RNA-seq was not generated in this study). Following LPS stimulation for 4 hours, both WT and HET macrophages underwent extensive transcriptional reprogramming (**Extended Data Fig. 1b**), yet the bias towards upregulation of H3K27me3-high genes remained evident (**Fig. 2g,h** and **Supplementary Table 11**). Approximately half of Cluster_1 DEGs upregulated in steady-state remained upregulated following LPS stimulation and were enriched for LPS-non-responsive genes (**Fig. 2i,j**), indicating that *DNMT3A* HET mutations confer a constitutive increase in expression that is maintained independently of inflammatory stimulation. These data indicate that *DNMT3A* mutations preferentially increase expression of bivalent Polycomb targets in macrophages, and that this signature is maintained in both resting and activated states.

### Baseline DNA methylation and H3K27me3 levels predict sensitivity to DNMT3A dysfunction

Having identified H3K27me3-high bivalent promoters as preferentially upregulated in *DNMT3A*-mutant macrophages, we next sought to understand the underlying mechanism. Because Polycomb-mediated repression involves chromatin compaction, we hypothesized that derepressed bivalent genes would be accompanied by increased promoter accessibility. ATAC-seq analysis revealed that H3K27me3-high Cluster_1 promoters display significantly greater accessibility in both HET and R882H mutant macrophages relative to WT, a phenotype markedly more pronounced than that observed at H3K27me3-low Cluster_2 promoters (**Fig. 3a**). These findings indicate that increased promoter accessibility accompanies derepression of bivalent Polycomb target genes in *DNMT3A*-mutant macrophages.

**Fig. 3.**
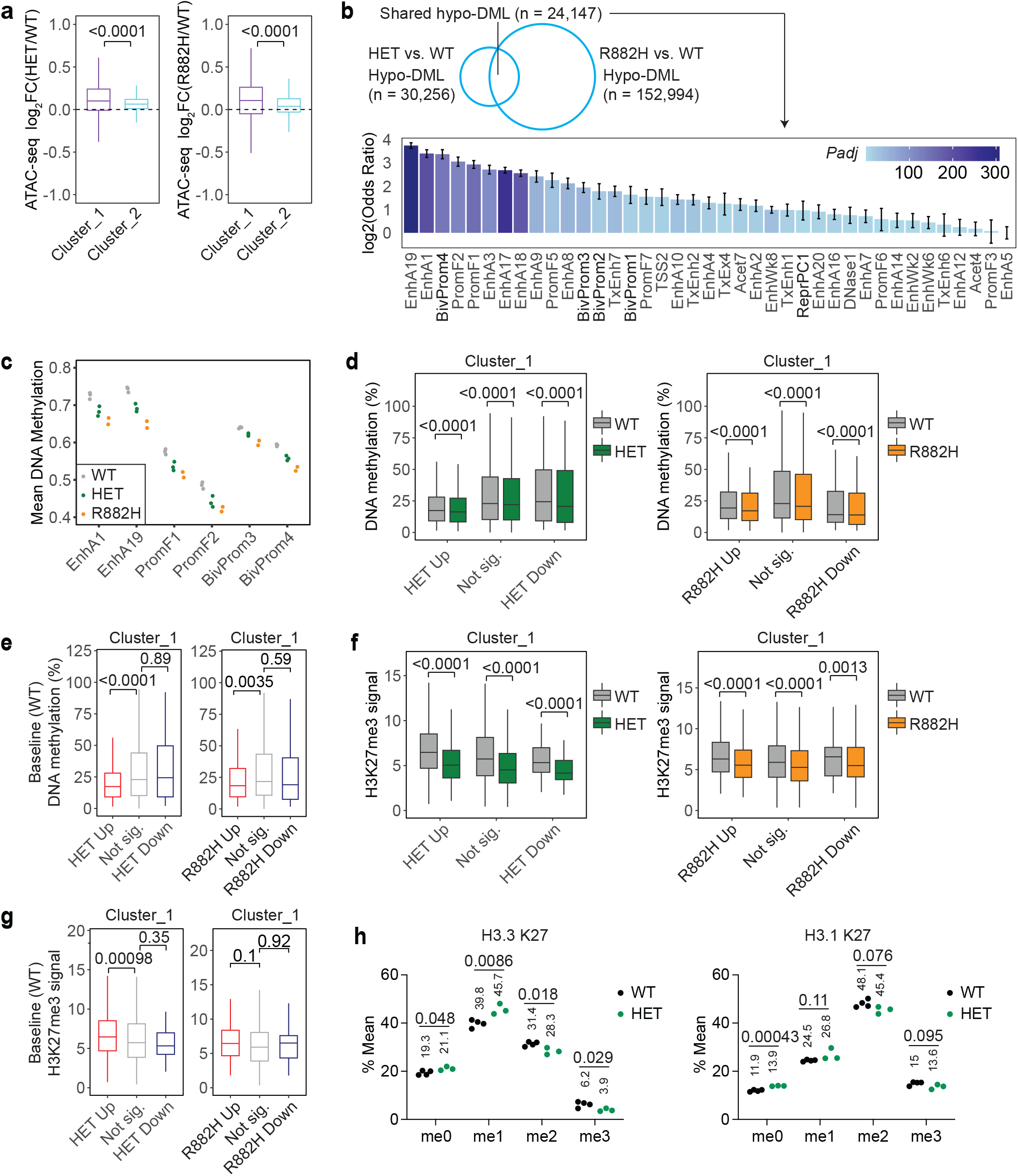
Baseline DNA methylation and H3K27me3 at bivalent promoters predict selective derepression in *DNMT3A*-mutant macrophages. **a,** ATAC-seq log2FC for HET versus WT or R882H versus WT macrophages grouped by H3K27me3-high bivalent genes (Cluster_1) and H3K27me3-low genes (Cluster_2), as defined in Fig. 2c. *P* values were calculated by Student’s *t*-test. **b,** Top: number of overlapping hypomethylated differentially methylated loci (hypo-DML; DSS DMLtest *P* < 0.01) between HET and R882H macrophages. Bottom: log2 odds ratios showing enrichment of hypo-DML shared between HET and R882H macrophages across a tissue-independent full-stack chromatin state annotation^34^. EnhA, active enhancer; BivProm, bivalent promoter; PromF, promoter flanking region; TxEnh, transcribed enhancer; TSS, transcription start site; TxEx, transcribed exon; Acet, acetylated region; EnhWk, weak enhancer; ReprPC, Polycomb-repressed. **c,** Mean methylation across CpGs within selected chromatin states shown in **b**. Each dot represents an independent clone, colored by genotype (WT, HET, R882H). **d,** Comparison of DNA methylation at Cluster_1 genes in WT versus *DNMT3A*-mutant macrophages, grouped by RNA-seq expression change. *P* values were calculated by a paired Wilcoxon signed-rank test, with pairing by CpG site. **e,** Baseline promoter DNA methylation in WT macrophages at Cluster_1 genes, grouped according to their RNA-seq expression changes in HET or R882H macrophages. *P* values were calculated by a Wilcoxon rank-sum test. **f,** Comparison of H3K27me3 levels measured by CUT&RUN at Cluster_1 genes in WT versus *DNMT3A*-mutant macrophages, grouped by RNA-seq expression change. *P* values were calculated by a paired Wilcoxon signed-rank test, with pairing by gene. **g,** Baseline promoter H3K27me3 levels in WT macrophages at Cluster_1 genes, grouped according to their RNA-seq expression changes in HET or R882H macrophages. *P* values were calculated by a Wilcoxon rank-sum test. **h,** Relative abundance of H3K27 methylation states (me0, me1, me2, and me3) on histone H3.3 and H3.1 in WT and HET macrophages; R882H clones were not analyzed. Quantified by mass spectrometry and expressed as a percentage of total H3.3 or H3.1 K27-containing peptide. Each dot represents an independent clone. *P* values were calculated by Student’s *t*-test.

To understand the contribution of DNA methylation in bivalent gene derepression, we performed genome-wide DNA methylation profiling by Enzymatic Methyl-Seq (EM-seq) and mapped the results onto a universal (i.e. cell-type independent) chromatin state annotation.^33,34^ Differentially methylated loci (DML) shared between HET and R882H macrophages were predominantly hypomethylated and strongly enriched in active enhancer (EnhA), promoter-flanking (PromF), and bivalent promoter (BivProm) chromatin states (**Supplementary Table 12**, **Fig. 3b**). In contrast, shared hypermethylated DML showed substantially weaker enrichment (maximum -log10 adjusted p ∼10, versus ∼300 for hypomethylated DML) (**Supplementary Table 13, Extended Data Fig. 2a**). Accordingly, average CpG methylation was reduced in EnhA, PromF, and BivProm regions in mutant macrophages (**Fig. 3c** and **Extended Data Fig. 2b)**. These results are consistent with previous reports of enhancer and Polycomb-target hypomethylation in *DNMT3A*-mutant hematopoietic cells.^13,19,21,23^

Notably, reduced DNA methylation at bivalent promoters was not restricted to upregulated genes (**Fig. 3d**). Rather, what distinguished upregulated genes was their baseline DNA methylation: in WT macrophages, promoters of upregulated Cluster_1 genes exhibited significantly lower methylation than those of unchanged genes (**Fig. 3e**). Studies in *Dnmt* triple-knockout or *Dnmt1*-knockout murine cells have demonstrated that bivalent genes undergo H3K27me3 depletion and gene derepression under conditions of profound DNA hypomethylation, with the most pronounced effects at genes carrying low baseline DNA methylation and high baseline H3K27me3.^35–37^ To test whether H3K27me3 is reduced in *DNMT3A*-mutant macrophages despite their much more modest degree of DNA methylation loss, we compared H3K27me3 levels between WT and mutant macrophages using CUT&RUN. H3K27me3 levels were reduced across all Cluster_1 gene subsets in *DNMT3A*-mutant macrophages, regardless of gene expression changes (**Fig. 3f**). At baseline, upregulated Cluster_1 genes showed significantly higher H3K27me3 in WT macrophages than unchanged Cluster_1 genes (**Fig. 3g**), consistent with stronger Polycomb activity at promoters that undergo derepression in *DNMT3A*-mutant macrophages.

Mass spectrometry analysis corroborated these locus-level findings at the global scale, revealing a moderate but significant reduction in H3K27me3 and H3K27me2 accompanied by reciprocal increases in unmethylated and mono-methylated H3K27 in HET macrophages (**Fig. 3h**). Notably, this effect was more pronounced on the histone variant H3.3, which is enriched at active genes and developmental loci, than on the canonical replication-coupled H3.1.^38–40^ The reduction in global H3K27me3 is consistent with findings in *Dnmt3a* R878H hematopoietic cells – the murine equivalent of the human R882H mutation.^41,42^ In contrast, other histone modifications examined, including H3K9 and H3K4 methylation, remained unchanged in HET macrophages (**Extended Data Fig. 2c**).

Taken together, these data indicate that across bivalent promoters, *DNMT3A* mutations broadly reduce DNA methylation and H3K27me3 and increase chromatin accessibility. Baseline chromatin state at these loci predicts sensitivity to DNMT3A dysfunction: genes with low promoter DNA methylation and high H3K27me3 at baseline are most vulnerable to transcriptional derepression.

### Bivalent Polycomb target gene derepression is independent of the developmental timing of *DNMT3A* mutation

We examined whether the preferential upregulation of bivalent Polycomb target genes observed in *DNMT3A*-mutant macrophages is already evident at earlier stages of hematopoietic differentiation. To address this, we sorted two progenitor populations along the *in vitro* hESC-to-macrophage differentiation trajectory: CD34+CD43+ and CD34−CD43+CD11b+ populations, corresponding to early hematopoietic progenitors and committed myeloid progenitors, respectively (**Extended Data Fig. 3a**). Bulk RNA-seq revealed that *DNMT3A*-mutant cells showed enrichment of PRC2-related gene sets (CGP GSEA) and development- and morphogenesis-related biological processes (GO GSEA) relative to WT controls in myeloid progenitors, but not in hematopoietic progenitors (**Extended Data Fig. 3b–e** and **Supplementary Table 14–19**), recapitulating the Polycomb-associated transcriptional signature observed in DNMT3A-mutant macrophages. However, the specific bivalent Polycomb target genes upregulated in *DNMT3A*-mutant macrophages were largely distinct from those upregulated in myeloid progenitors, with only limited overlap between the two populations (**Extended Data Fig. 3f**). Together, these findings indicate that derepression of Polycomb target genes begins at or before the myeloid progenitor stage, but that the specific transcriptional response at individual bivalent loci might be differentiation-stage dependent.

While *DNMT3A* mutations in clonal hematopoiesis are thought to arise in hematopoietic stem and progenitor cells,^43,44^ our hESC models carry *DNMT3A* mutations from the pluripotent stem cell stage. We therefore asked whether the developmental timing of *DNMT3A* mutation acquisition influences Polycomb target gene derepression in differentiated macrophages. We studied macrophages derived from mice carrying a Cre-inducible R878H mutation in the endogenous *Dnmt3a* locus,^45^ crossed with *Vav*-Cre mice to induce the mutation specifically within the hematopoietic compartment (*Dnmt3a^fl-^*^R878H/+^;*Vav*-Cre, hereafter Dnmt3a-R878H).

Bone marrow cells from WT or Dnmt3a-R878H mice were differentiated *in vitro* into bone marrow-derived macrophages (BMDMs) (**Fig. 4a,b**).

**Fig. 4.**
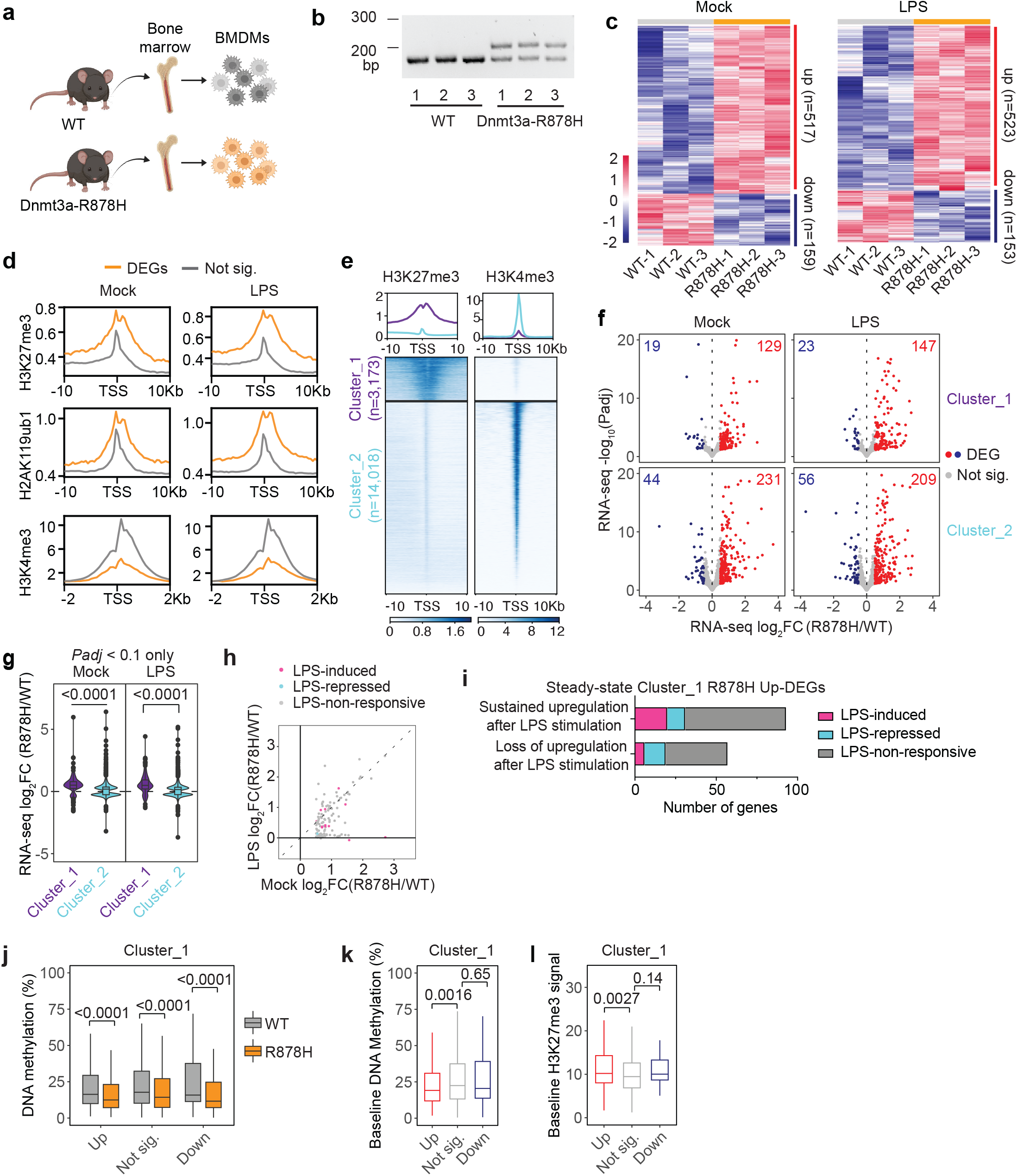
Bivalent Polycomb target genes are derepressed in Dnmt3a-R878H bone marrow– derived macrophages. **a,** Schematic of the experimental design using mouse bone marrow–derived macrophages (BMDMs) differentiated *in vitro* from bone marrow of WT or Dnmt3a^fl-R878H/+^; Vav-Cre (Dnmt3a- R878H) mice.^45^ **b**, PCR genotyping of genomic DNA from BMDMs of the three WT and three Dnmt3a-R878H mice used for the analyses shown in **c**–**l**. Primers yield products of 178 bp (WT) and 282 bp (R878H). **c,** Heatmap of DEGs (Padj < 0.1 and |log2FC| > 0.5) between WT and Dnmt3a-R878H BMDMs under mock (PBS) or LPS-stimulated conditions. Rows represent DEGs and columns represent individual mice; colors indicate row-scaled expression (z-score; red, higher expression; blue, lower expression). **d,** Distribution of H3K27me3, H2AK119ub, and H3K4me3 signal at TSSs and surrounding regions for Dnmt3a-R878H DEGs (Padj < 0.1 and |log2FC| > 0.5) versus non-significant genes (Not sig.), based on CUT&RUN data from WT BMDMs. **e,** Heatmaps of H3K27me3 and H3K4me3 signal within a 20-kb window centered on the TSSs of expressed genes in WT BMDMs, partitioned by k-means clustering (k = 2) on H3K27me3 signal using deepTools to define H3K27me3-high promoters (Cluster_1) and H3K27me3-low promoters (Cluster_2). The number of genes in each cluster is indicated. **f,** Volcano plots of shrunken log2FC (Dnmt3a-R878H versus WT) against −log10(Padj) under mock and LPS-stimulated conditions (100 ng/ml, 4 h). Colored dots indicate DEGs (Padj < 0.1 and |log2FC| > 0.5; red, upregulated; blue, downregulated); gray dots indicate non-DEGs. The numbers of upregulated and downregulated DEGs are indicated. **g,** Violin plots of shrunken log2FC (Dnmt3a-R878H versus WT) for genes with Padj < 0.1 in Cluster_1 and Cluster_2 under mock and LPS-stimulated conditions, illustrating the bias toward upregulation among Cluster_1 genes. P values were calculated by a Wilcoxon rank-sum test. **h**, Relationship between Dnmt3a-R878H versus WT shrunken log2FC under mock and LPS-stimulated conditions for Cluster_1 genes upregulated in Dnmt3a-R878H mBMDMs under mock conditions (Padj < 0.1 and log2FC > 0.5). Dots are colored according to transcriptional response to LPS: LPS-induced, LPS versus mock in WT mBMDMs, Padj < 0.1 and log2FC > 1; LPS-repressed, Padj < 0.1 and log2FC < −1; LPS-non-responsive, all remaining genes. **i**, Cluster_1 genes upregulated in Dnmt3a-R878H versus WT mBMDMs under mock conditions, as defined in **h**, grouped according to whether they remained upregulated or were no longer differentially expressed under LPS-stimulated conditions. Each group is further classified based on transcriptional response to LPS as LPS-induced, LPS-repressed, or LPS-non-responsive, as defined in **h**. **j**-**l**. Boxes show the median and interquartile range (IQR); whiskers extend to 1.5 × IQR. **j,** DNA methylation levels measured by EM-seq at promoters of Cluster_1 genes in WT versus Dnmt3a-R878H BMDMs, grouped by RNA-seq expression change in Dnmt3a-R878H versus WT (Up, Not sig., Down). P values were calculated by a paired Wilcoxon signed-rank test, with pairing by CpG site. **k,** Baseline promoter DNA methylation in WT BMDMs at Cluster_1 genes, grouped according to their RNA-seq expression change in Dnmt3a-R878H BMDMs. P values were calculated by a Wilcoxon rank-sum test. **l,** Baseline promoter H3K27me3 levels measured by CUT&RUN in WT BMDMs at Cluster_1 genes, grouped according to their RNA-seq expression change in Dnmt3a-R878H BMDMs. P values were calculated by a Wilcoxon rank-sum test.

Bulk RNA-seq revealed significant gene expression changes in Dnmt3a*-*R878H macrophages relative to WT in both steady-state and LPS-stimulated conditions (**Fig. 4c** and **Supplementary Table 20**). To determine whether bivalent Polycomb target genes are similarly affected in this hematopoietic lineage-restricted model, we overlaid DEGs with H3K27me3, H2AK119ub1, and H3K4me3 profiles generated in-house from WT BMDMs. As in hESC-derived macrophages, DEGs in Dnmt3a-R878H BMDMs were enriched for promoters marked by H3K27me3 and H2AK119ub1 relative to non-DEGs (**Fig. 4d**). Using the WT BMDM histone profiles, we clustered expressed genes based on promoter H3K27me3 levels to define H3K27me3-high genes (Cluster_1) and H3K27me3-low genes (Cluster_2) (**Fig. 4e** and **Supplementary Table 21**). Within Cluster_1, DEGs were more likely to be upregulated than downregulated in Dnmt3a-R878H BMDMs in both resting and LPS-stimulated states (**Fig. 4f,g**), mirroring the bias towards derepression of bivalent Polycomb targets observed in *DNMT3A*-mutant hESC-derived macrophages.

As in the human system, the majority of Cluster_1 DEGs upregulated at steady state in Dnmt3a-R878H BMDMs remained upregulated following LPS stimulation and were enriched for LPS-non-responsive genes (**Fig. 4h,i**), indicating that the constitutive, stimulation-independent increase in expression conferred by *DNMT3A* mutation is conserved between species.

Together, these data indicate that upregulation of bivalent Polycomb target genes is a consistent feature of macrophages differentiated from *DNMT3A*-mutant precursors, regardless of whether the mutation is acquired at the pluripotent stem cell stage or in the hematopoietic compartment, and that this constitutive derepression persists independently of inflammatory stimulation in both human and murine macrophages.

We next assessed the relationship between DNA methylation and gene expression in this murine system. EM-seq of WT and Dnmt3a-R878H BMDMs showed that DNA methylation was reduced across Cluster_1 promoters in Dnmt3a-R878H macrophages regardless of gene expression changes (**Fig. 4j**), recapitulating the pattern observed in *DNMT3A*-mutant hESC-derived macrophages. In WT BMDMs, promoters of upregulated Cluster_1 genes exhibited significantly lower baseline DNA methylation and higher baseline H3K27me3 than promoters of unchanged Cluster_1 genes (**Fig. 4k,l**), analogous to the human system. These findings indicate that the preferential derepression of bivalent Polycomb target genes with low baseline promoter methylation and high H3K27me3 is conserved across species and across developmental contexts in which *DNMT3A* mutations are acquired.

### Enhanced macrophage migration and injury response are associated with derepression of bivalent Polycomb target genes

We hypothesized that derepression of bivalent Polycomb target genes might alter macrophage behavior. To investigate this, we examined biological pathways jointly upregulated at steady state across four macrophage datasets: our in-house RNA-seq from human DNMT3A-HET and DNMT3A-R882H hESC-derived macrophages and murine Dnmt3a-R878H BMDMs, together with publicly available RNA-seq from murine BMDMs with germline Dnmt3a haploinsufficiency (Dnmt3a-HET).^18^ Gene Ontology (GO) analysis of genes upregulated across these fou datasets identified several consistently enriched pathways (**Supplementary Table 22–24**).^46^ Integration with H3K27me3 profiles revealed that the majority of these jointly enriched pathways were dominated by Polycomb target genes, including actin-filament based process (GO:0030029), extracellular matrix organization (GO:0030198), and response to wounding (GO:0009611) (**Fig. 5a** and **Extended Data Fig. 4a**). In contrast, inflammation-related pathways, such as inflammatory response (GO:0006954) and positive regulation of cytokine production (GO:0001819), were not enriched for Polycomb target genes (**Fig. 5a** and **Extended Data Fig. 4a**). This suggests that, although *DNMT3A*-mutant macrophages show enhanced inflammatory gene expression as reported previously,^12–18^ these effects are not a direct consequence of Polycomb target gene derepression.

**Fig. 5.**
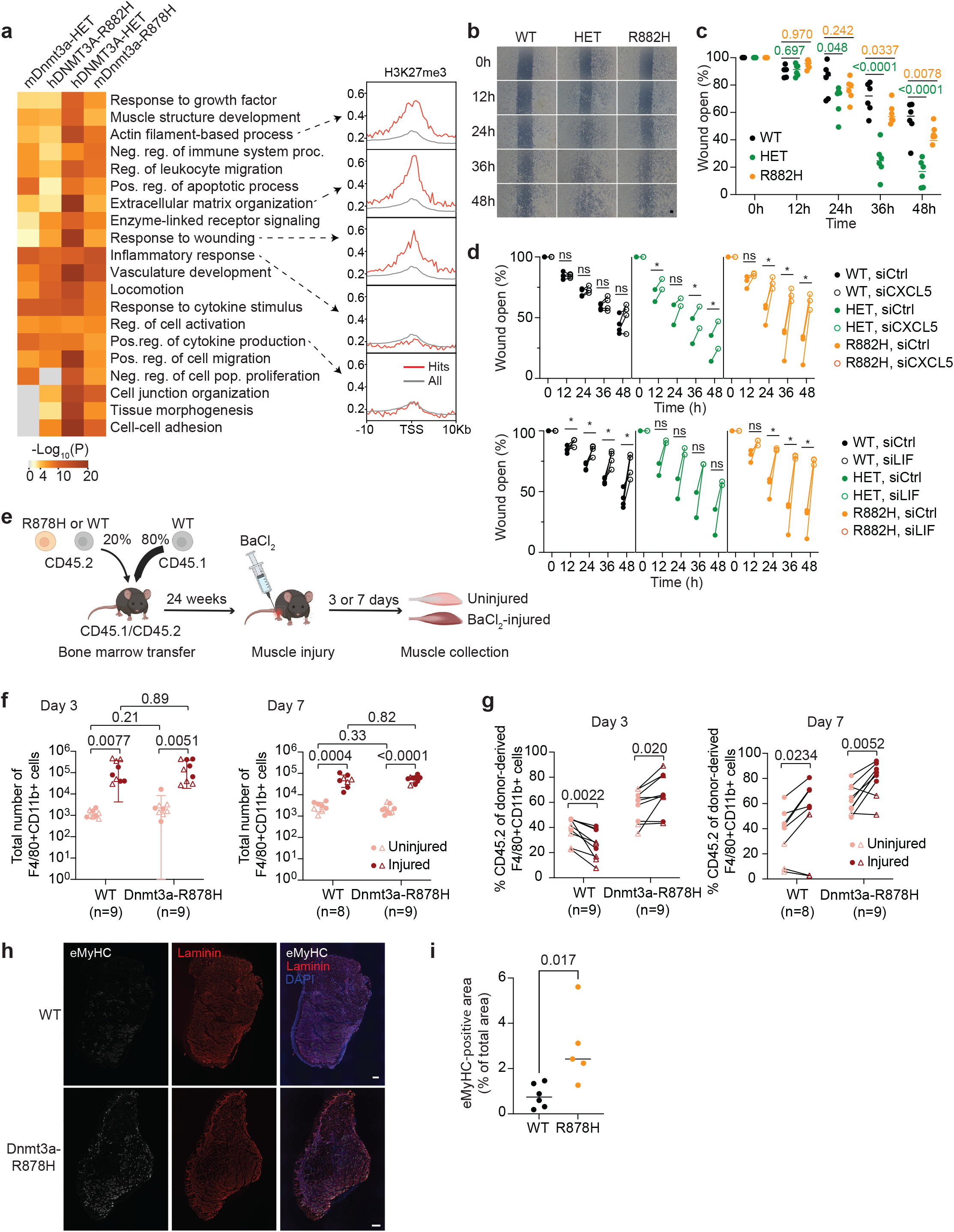
*DNMT3A*-mutant macrophages exhibit enhanced migratory capacity *in vitro* and *in vivo*. **a,** Left: heatmap of the top 20 enriched Gene Ontology (GO) terms for four sets of upregulated DEGs in human and murine macrophages (mDnmt3a-HET: murine Dnmt3a-HET^18^; hDNMT3A-R882H: human DNMT3A-R882H; hDNMT3A-HET: human DNMT3A-HET; and mDnmt3a-R878H: murine Dnmt3a-R878H), colored by enrichment *P* value (Metascape^46^). Right: H3K27me3 enrichment analysis for selected GO terms; “Hits” denotes DEGs within each GO term and “All” denotes all expressed genes. **b,c,** Wound healing assay of WT, HET, and R882H human macrophages. **b**, Representative phase-contrast images of one WT, one HET, and one R882H clone at the indicated time points after scratch injury. Scale bar, 200 µm. **c**, Quantification of remaining wound area over time for three WT, three HET, and three R882H clones. *P* values were calculated by two-way ANOVA (genotype × time). **d,** Quantification of remaining wound area over time in WT, HET and R882H macrophages following siRNA-mediated knockdown of *CXCL5* or *LIF* for 48 h. siCtrl, non-targeting siRNA. *P* values were calculated by a paired t-test (siRNA versus siCtrl within each clone) and are indicated as *, *P* ≤ 0.1; ns, not significant. **e**, Experimental design for *in vivo* recruitment assays. Mixed bone marrow chimeras were generated by transplanting a 1:4 (test:competitor) mixture of WT or Dnmt3a-R878H test bone marrow (CD45.2) and WT competitor bone marrow (CD45.1) into lethally irradiated WT recipients (CD45.1/CD45.2). Twenty-four weeks after transplantation, recipients received BaCl_2_-induced injury to the quadriceps and tibialis anterior (TA) muscles of one hindlimb, with the contralateral hindlimb left uninjured as an internal control. Quadriceps muscles were used for flow cytometry (f, g) and TA muscles for immunofluorescence (h, i). **f,g**, Flow cytometric quantification of myeloid cell recruitment to quadriceps muscle. The number of mice analyzed per group is indicated. Filled circles denote female mice; open triangles denote male mice. **f,** Total number of myeloid cells (F4/80^+^CD11b^+^) in uninjured and injured quadriceps muscle at day 3 and day 7 post-injury. Within each mouse, injured and contralateral uninjured limbs were compared by a paired t-test; injured limbs were compared between WT and Dnmt3a-R878H chimeras by Welch’s t-test. **g,** Percentage of test-derived (CD45.2^+^) cells among F4/80^+^CD11b^+^ myeloid cells in quadriceps muscle at day 3 and day 7 post-injury. For each mouse, the injured limb was compared with the contralateral uninjured limb, which serves as the within-animal baseline for engraftment. *P* values were calculated by a paired t-test (injured versus uninjured within each mouse). **h**, Representative images of TA muscle stained for embryonic myosin heavy chain (eMyHC), laminin, and DAPI at day 7 post-injury. Remaining muscle sections are shown in Extended Data Fig. 5c. Scale bar, 200 µm. **i**, Quantification of eMyHC-positive area as a percentage of total area (n = 6 WT and n = 5 Dnmt3a-R878H mice) for TA muscle at day 7 post-injury. Muscles with processing artifacts precluding reliable quantification were excluded. *P* values were calculated by a Mann-Whitney test.

Notably, many of the H3K27me3-enriched pathways were related to cell migration and wound repair. To determine whether this transcriptional signature correlates with functional changes, we performed scratch assays using WT, HET and R882H hESC-derived macrophages. Both HET and R882H macrophages exhibited significantly faster wound closure than WT macrophages, although this effect was more pronounced in HET macrophages (**Fig. 5b, c**). This occurred without evidence of increased proliferation (**Extended Data Fig. 4b**), indicating that the faster wound closure reflects increased migratory capacity rather than proliferation. The more pronounced migratory phenotype in HET macrophages paralleled the stronger transcriptional enrichment of cell migration and wound repair-related pathways in HET relative to R882H macrophages (**Fig. 5a**).

To identify bivalent Polycomb target genes underlying the enhanced migratory phenotype, we selected candidates that were H3K27me3-high, upregulated in both HET and R882H hESC-derived macrophages (adjusted p < 0.1, log2FC >2), and had established links to cell migration. These candidates were further narrowed to two by qPCR analysis of RNA from independent batches of macrophages (**Extended Data Fig. 4c**). siRNA-mediated knockdown (**Extended Data Fig. 4d**) showed that LIF depletion delayed wound closure in both WT and mutant macrophages, whereas depletion of CXCL5 significantly delayed wound closure only in *DNMT3A*-mutant macrophages (**Fig. 5d**), indicating that LIF supports migration broadly across genotypes whereas CXCL5 acts selectively in the DNMT3A mutant context.

We next asked whether ongoing Polycomb activity is required to maintain this phenotype once macrophages have terminally differentiated, or whether the underlying epigenetic changes are established earlier and no longer depend on continuous Polycomb function in mature cells. To distinguish between these possibilities, we performed siRNA-mediated knockdown of RNF2 and EZH2, core components of PRC1 and PRC2, respectively, in terminally differentiated macrophages (**Extended Data Fig. 4e**). Acute knockdown of EZH2 or RNF2 did not alter migration under the conditions tested (**Extended Data Fig. 4f**), indicating that ongoing PRC1/PRC2 activity is not required to maintain the migratory phenotype in mature macrophages, and suggesting instead that the relevant epigenetic changes are established prior to terminal macrophage differentiation.

We next asked whether *Dnmt3a*-mutant myeloid cells show altered recruitment to sites of tissue injury *in vivo*. To mimic clonal hematopoiesis, we generated mixed bone marrow chimeras by transplanting a 1:4 mixture of WT or Dnmt3a-R878H donor bone marrow (CD45.2) with WT competitor bone marrow (CD45.1) into lethally irradiated WT recipients (CD45.1/CD45.2 heterozygotes), enabling discrimination of donor-derived cells by CD45 congenic markers (**Fig. 5e**). Consistent with previous studies,^45,47^ Dnmt3a-R878H donor cells exhibited a competitive advantage: by day 37 post-transplant, CD45.2^+^ cells already constituted a significantly larger fraction of peripheral blood leukocytes in Dnmt3a-R878H chimeras than in WT chimeras, and this difference became even more pronounced over time (**Extended Data Fig. 5a** and **Supplementary Table 25**). After 24 weeks of reconstitution, we induced acute muscle injury by injecting BaCl2 into the quadriceps and tibialis anterior of one hindlimb, leaving the contralateral hindlimb uninjured as an internal control. In this model, immune cell infiltration peaks within the first few days after injury and is followed by a regenerative phase by approximately day 7.^48^ At 3 and 7 days post-injury, quadriceps were harvested for flow cytometric analysis of immune cell infiltration (**Extended Data Fig. 5b**) and tibialis anterior muscles were processed for histology.

Injured quadriceps contained substantially more myeloid cells than uninjured quadriceps at both time points (**Fig. 5f** and **Supplementary Table 26**). At day 3, donor-derived CD45.2⁺ cells made up a higher fraction of the myeloid compartment in injured than in contralateral uninjured quadriceps in Dnmt3a-R878H chimeras, whereas the reverse was true in WT chimeras (**Fig. 5g** and **Supplementary Table 26**). By day 7, donor fractions were higher in injured quadriceps in both genotypes (**Fig. 5g** and **Supplementary Table 26**). Mutant donor cells were therefore enriched at the injury site at both time points, whereas WT donor cells became enriched only during the regenerative phase, consistent with earlier recruitment of mutant myeloid cells. This occurred without differences in total myeloid numbers between genotypes (**Fig. 5f** and **Supplementary Table 26**), indicating a shift in the composition rather than the magnitude of the infiltrate.

To assess whether altered myeloid recruitment is accompanied by differences in tissue repair, we stained injured tibialis anterior muscles at day 7 post-injury for embryonic myosin heavy chain (eMyHC), a marker of newly formed regenerating myofibers. The eMyHC-positive area, expressed as a percentage of total tissue area, was higher in Dnmt3a-R878H than in WT chimeras (**Fig. 5h,i, Extended Data Fig. 5c** and **Supplementary Table 27)**. Together with the enhanced migratory capacity observed in scratch assays, these findings indicate that *DNMT3A*-mutant myeloid cells exhibit a selective advantage in early recruitment to sites of tissue injury and are associated with increased myofiber regeneration, consistent with derepression of bivalent Polycomb target genes involved in cell migration and wound response.

## Discussion

Somatic *DNMT3A* mutations are the most common lesions in clonal hematopoiesis,^3,4^ and produce well-described DNA methylation changes in hematopoietic stem and progenitor cells.^1,19,21,49,50^ Polycomb-associated regulatory regions are known preferential targets of this methylation loss; notably, Nam et al. showed that DNMT3A-R882H human CD34⁺ progenitors display preferential hypomethylation at PRC2-associated motifs and bivalent (H3K27me3– H3K4me3) regions, but did not observe corresponding gene expression changes at PRC2 target genes.^21^ This raises the question of whether, and under what circumstances, hypomethylation of Polycomb-associated regions is translated into transcriptional derepression. Here, using isogenic human and murine macrophage models with multi-layered epigenomic profiling, we address this question directly and link it to macrophage function. We show that DNMT3A mutations broadly reduce H3K27me3 and increase chromatin accessibility at bivalent promoters, alongside the expected DNA hypomethylation, but that only a selective subset of bivalent genes is derepressed. This relationship is conserved between human and murine macrophages, does not require DNMT3A mutation from the pluripotent stage, and is accompanied by enhanced macrophage migration and preferential recruitment to injured tissue.

In murine *Dnmt3a*-mutant HSCs, hypomethylation and transcriptional change are frequently uncoupled, with many methylation changes occurring without corresponding alterations in gene expression.^1,19,51^ Our data extend this observation to macrophages and identify the chromatin feature that distinguishes responsive from unresponsive loci. DNA methylation and H3K27me3 at bivalent promoters were reduced in mutant macrophages regardless of expression change (**Fig. 3d,f**, **Fig. 4j**), indicating that methylation loss alone is not sufficient to drive transcriptional change and that macrophages are not a uniquely sensitive lineage. What distinguished responsive loci was their pre-existing configuration: promoters that became derepressed already carried lower DNA methylation and higher H3K27me3 in WT cells than those that remained unchanged (**Fig. 3e,g**, **Fig. 4k,l**), indicating that the transcriptional consequences of DNMT3A loss depend on the chromatin context—offering one explanation for why Polycomb-target hypomethylation does not uniformly result in derepression. Consistent with this reflecting a change in resting chromatin state rather than signal-responsiveness, elevated expression of bivalent targets was largely maintained after LPS stimulation and was most apparent among genes that are themselves not LPS-responsive (**Fig. 2i,j**, **Fig. 4h,i**), indicating that *DNMT3A* mutations principally reset baseline expression rather than altering inducibility.

The derepressed promoters share a configuration that dovetails with the known targeting of DNMT3A1, the predominant isoform in differentiated cells, to Polycomb-regulated regions via N-terminal recognition of PRC1-deposited H2AK119ub.^52,53^ Together, these observations support a model in which DNMT3A is recruited to promoters with high Polycomb occupancy and contributes to the maintenance of the surrounding Polycomb chromatin state. Consistent with this, promoters of DNMT3A-responsive genes were enriched for H2AK119ub as well as H3K27me3 in both human and murine macrophages (**Fig. 2b**, **Fig. 4d**). Because macrophages lack the sharply bounded H3K27me3 domains characteristic of ESCs, our CUT&RUN data cannot distinguish boundary erosion from reduced within-domain density; nonetheless, the results are compatible with models in which DNA methylation contributes to the integrity of Polycomb domains.^36^ Definitive resolution will require separation-of-function DNMT3A alleles that disrupt PRC-directed targeting without globally abolishing de novo methylation.

Because our hESC models carry *DNMT3A* mutations from the pluripotent state, whereas clonal hematopoiesis mutations arise in HSPCs, we addressed the role of developmental timing from two directions. Along the hESC-to-macrophage trajectory, PRC2 and developmental gene set enrichment was not detected in early hematopoietic progenitors but became evident in committed myeloid progenitors, indicating that Polycomb target derepression is established before terminal macrophage differentiation. Complementarily, macrophages differentiated from mice in which Dnmt3a-R878H was induced specifically in the hematopoietic compartment reproduced the same molecular features. Thus, preferential derepression of bivalent Polycomb targets is a robust consequence of DNMT3A dysfunction that does not require the mutation to be present from the pluripotent stage and is conserved across human and murine macrophages. Notably, the loci upregulated in myeloid progenitors showed limited overlap with those upregulated in macrophages, raising the possibility that although the effects of DNMT3A dysfunction emerge during early hematopoietic differentiation, the specific loci that become derepressed may be shaped by the chromatin and transcriptional context of each developmental stage.

Haploinsufficiency and the R882H hotspot allele have been proposed to have distinct molecular mechanisms, with R882H reported to exert dominant-negative effects and to display altered flanking sequence preferences.^24,54–56^ We therefore analyzed them as separate genotypes throughout. Both converge on the same qualitative phenotype—enrichment of bivalent chromatin among DEGs, biased upregulation of H3K27me3-high genes, increased promoter accessibility, and accelerated wound closure—supporting reduced DNMT3A function at Polycomb-occupied promoters as a shared consequence of these mutations. However, the two alleles are not equivalent: the PRC2-associated signature was more pronounced in HET macrophages, whereas the inflammatory signature was stronger in R882H (**Fig. 5a**). This dissociation is consistent with our broader finding that the inflammatory transcriptional signature of *DNMT3A*-mutant macrophages is not directly attributable to bivalent Polycomb derepression, since inflammatory pathways were not enriched for Polycomb targets (**Fig. 5a**) and a substantial fraction of DEGs in both species belong to Cluster_2 (low H3K27me3) (**Fig. 2e**, **Fig. 4f**). Such changes may instead reflect indirect consequences of derepression or independent mechanisms such as enhancer hypomethylation, the most prominent methylation change in our EM-seq data (**Fig. 3b**).

Functionally, the derepressed Polycomb program is enriched for genes governing migration, extracellular matrix remodeling, and wound response, and *DNMT3A*-mutant macrophages of both genotypes closed scratch wounds faster than wild-type without evidence of increased proliferation. Targeted knockdown of individual derepressed bivalent targets confirmed that genes within this program can contribute to migratory behavior, one acting broadly across genotypes and another selectively in the mutant context. We regard these as illustrative rather than exhaustive: hundreds of bivalent loci are derepressed and their individual expression changes are modest, so the phenotype most likely reflects the combined effect of many such changes rather than a single dominant effector. *In vivo*, mutant myeloid cells were preferentially recruited to injured muscle during the early inflammatory phase without a change in total myeloid burden, indicating a shift in the composition rather than the magnitude of the infiltrate, and this was accompanied by a greater area of regenerating myofibers. Together, these findings link *DNMT3A* mutation-associated Polycomb derepression to altered macrophage migration and preferential recruitment to injured tissue, with potential consequences for tissue repair.

Several limitations of our study warrant mention. Our models are macrophages differentiated *ex vivo*, which afford mechanistic control but may not capture the cues that shape macrophages *in vivo*; whether comparable bivalent promoter derepression occurs in macrophages in individuals with clonal hematopoiesis remains to be determined. Our chromatin measurements are correlative, and although the association between baseline promoter state and derepression is consistent across species, we have not established that H3K27me3 loss is causal. Finally, while individual derepressed targets contribute to macrophage migration *in vitro*, we have not identified which targets are necessary or sufficient for the recruitment and regeneration phenotypes observed *in vivo*, nor whether the altered regenerative dynamics we observe represent accelerated repair, a shift in the timing of inflammatory resolution, or a change that becomes maladaptive under chronic or repeated injury.

In summary, we find that *DNMT3A* mutations are associated with altered bivalent Polycomb chromatin in macrophages and that the resulting transcriptional derepression is selective and predictable from baseline promoter DNA methylation and H3K27me3. This program is shared between human and murine macrophages, does not require the mutation to be acquired at the pluripotent stage, and is associated with enhanced macrophage migration, preferential early recruitment to injured tissue, and increased early myofiber regeneration in vivo. By linking baseline promoter chromatin features to susceptibility to derepression and connecting these molecular changes to macrophage behavior, this work provides a framework for understanding how somatic epigenetic lesions arising in stem cells can have functional consequences in differentiated innate immune cells.

## Methods

### hESC culture and maintenance

All experiments involving hESCs were approved by the UCI Human Stem Cell Research Oversight Committee (hSCRO). H1 (WA01) hESCs (WiCell) were maintained under feeder-free conditions in StemFlex medium (Cat# A3349401, Gibco, Thermo Fisher Scientific) on Cultrex RGF BME–coated plates (Cat# 3433-010-01, R&D Systems). Cells were passaged using Accutase (STEMCELL Technologies) and replated in StemFlex medium supplemented with 2 μM thiazovivin (Cat # S1459, Selleck Chemicals).

### Genome editing of hESCs

Genome editing was performed using Cas9 ribonucleoprotein (RNP) delivery. hESCs were resuspended in P3 nucleofector solution (Cat# V4XP-3032, Lonza) and nucleofected with pre-assembled RNPs comprising crRNA:tracrRNA duplexes and Alt-R HiFi Cas9 nuclease (Integrated DNA Technologies) using a 4D-Nucleofector X Unit (Lonza, Cat# AAF-1002B; program DN-100). Generation of the *DNMT3A*-heterozygous line was described previously^13^. For introduction of the *DNMT3A*-R882H mutation, single-stranded oligodeoxynucleotides (ssODNs) were included as homology-directed repair templates. Following nucleofection, cells were plated onto Cultrex-coated plates in StemFlex medium with thiazovivin and then transferred onto irradiated mouse embryonic fibroblasts (iMEFs, CF1 strain) to allow single-colony formation. Individual colonies were manually picked, expanded under feeder-free conditions, and genotyped by PCR amplification of CRISPR-targeted regions followed by Sanger sequencing. Colonies carrying the intended edit underwent one additional round of single-colony isolation to ensure clonality.

### Macrophage differentiation

hESC-derived macrophages were generated as previously described^13^ with minor modifications. For embryoid body (EB) formation, cells were resuspended at 1.25*10^5^ cells/ml in EB medium consisting of StemFlex supplemented with 50 ng/ml BMP-4 (Cat # 314-BP, R&D systems), 20 ng/ml stem cell factor (Cat # 255-SC, R&D systems), 50 ng/ml vascular endothelial growth factor (Cat # 293-VE, R&D systems), and 10 μM thiazovivin, and plated into 96-well ultra–low-attachment plates (Cat# 7007, Corning) for 8 days. After 8 days of EB differentiation, EBs were transferred to macrophage differentiation medium consisting of X-VIVO 15 (Cat # 02-053Q, Lonza) supplemented with 100 ng/ml M-CSF (Cat # 216-MC, R&D systems), 25 ng/ml IL-3 (Cat # 203-IL, R&D system), 2 mM GlutaMAX (Cat # 35050079, Gibco), 1% penicillin/streptomycin (Gibco), and 55 μM β-mercaptoethanol (Cat # 21985023, Gibco). Nonadherent macrophage precursors shed into the supernatant were collected approximately every 5 days and transferred to new tissue culture plates in macrophage maturation medium consisting of X-VIVO 15 supplemented with 50 ng/ml M-CSF. Cells were then maintained for >2 weeks in maturation medium to allow terminal differentiation. During the initial phase, macrophage precursors continued to divide, but proliferation progressively diminished by ∼2 weeks as cells acquired a terminally differentiated, cell cycle–arrested macrophage phenotype. Macrophages at this stage were used for transcriptomic, epigenomic and functional (scratch-wound) assays.

For experiments involving progenitor populations, cells were harvested at differentiation days 14 and 20 and stained with fixable viability dye eFluor 450 (eBioscience, Cat #65-0863-14) for 30 min at 4 °C. Fc receptors were blocked with FcR Blocking Reagent (Miltenyi Biotec, Cat #130-059-901) for 10 min at 4 °C, and cells were then stained with fluorochrome-conjugated antibodies against CD34 (BioLegend, Cat #343616), CD43 (BioLegend, Cat #343204), and CD11b (eBioscience, Cat #17-0118-42) at 4 °C protected from light. Cells were washed, resuspended in flow cytometry buffer (2% FBS in DPBS), and filtered through a 40 μm strainer before sorting on a BD FACSAria Fusion cell sorter. After gating on live singlets, early hematopoietic progenitors (CD34⁺CD43⁺) were sorted at day 14 and committed myeloid progenitors (CD34⁻CD43⁺CD11b⁺) at day 20. Sorted cells were collected directly into RNA lysis buffer (Zymo Research) and processed immediately for RNA isolation. Sorting was performed on four WT, two HET, and two R882H independent hESC clones.

### scRNA-seq

scRNA-seq libraries were generated with the 10x Genomic system Chromium Single Cell 3’ Kit (V3) following the manufacturer’s instructions and sequenced on an Illumina HiSeq 4000 instrument. Single-cells were de-multiplexed using Seurat’s (version 5.0.2)^57^ vignette for demultiplexing with hashtag oligos (https://satijalab.org/seurat/articles/hashing_vignette.html), as well as doublet identification, using the *HTODemux* function. Upon visual inspection of HTO signal distributions and cells in tSNE space, we observed that, for two HET samples (both with HTO.5 hashtag), ∼860-1,200 cells may have been mis-classified as doublets due to sequencer contamination with HTO.6, thus over-classifying HTO.5 cells as doublets. To recover cells for these two samples, we re-assigned doublet annotations by analyzing the correlation between the cells’ HTO.5 and HTO.6 signals (X-axis HTO.5 signal, Y-axis HTO.6 signal). Cells with signals deviating more than 1.5 standard deviations from the main regression line representing the majority population, were excluded. Additional doublets, identified as micro-clusters in the tSNE space derived from HTO signals, were removed. Cells initially classified as “Negative” (i.e., exhibiting low HTO.5 and HTO.6 signals) by the *HTODemux* function were also filtered out. For the remaining samples, we retained only cells assigned as “Singlets” by the *HTODemux* function. In total, 18,430 cells remained for downstream analyses.

We then performed the standard processing steps in *Seurat*. We first removed potential low-quality or stressed cells with percent mitochondrial reads exceeding three MAD above the median [median(percent.mt) + (3 x MAD)]. Raw counts for the remaining cells were then normalized using *NormalizeData*, followed by feature selection using *FindVariableFeatures*. The data was then scaled using *ScaleData*. For variance stabilization and normalization, we applied *SCTransform* with the parameter return.only.var.genes = F, allowing all genes to be retained for downstream analyses. Principal component analysis (PCA) was performed using *RunPCA*. We performed t-distributed stochastic neighbor embedding (t-SNE) using *RunTSNE* using the first 30 principal components. For cell clustering, we constructed a shared nearest neighbor (SNN) graph with *FindNeighbors* using the first 30 principal components and identified clusters using *FindClusters* with a resolution of 0.08. Differential expression analysis between WT and mutants was conducted using *FindMarkers* with parameters min.pct = 0.01, logfc.threshold = 0.01, and test.use = “MAST”. Gene set enrichment analysis (GSEA) was performed using the *GSEA* function in clusterProfiler^58^ version 4.10.1 on the MSigDB C2 CGP and GO:BP collection, obtained from the *msigdbr*^59^ package version 7.5.1 in R, with genes ranked by average log2 fold-change. Gene sets were considered enriched or depleted if adjusted p-value (Benjamini-Hochberg) < 0.05. PRC Module Scores were computed using the AddModuleScore_UCell function in UCell^60^. All commands were executed using default settings unless indicated otherwise.

### RNA-seq

RNA-seq was performed on hESC-derived macrophages, sorted hESC-derived progenitor populations, and mouse BMDMs. Total RNA was extracted using the Quick-RNA kit (Cat #R1051, Zymo Research), and RNA quality was assessed by RIN on an Agilent 2100 Bioanalyzer (Agilent Technologies). QC-passed libraries were prepared with the SMARTer mRNA-seq Library Prep Kit and sequenced on an Illumina NovaSeq 6000 as 100-bp paired-end reads. Reads were quantified with Salmon^61^ against a decoy-aware index built from GENCODE v.41, GRCh38, for human samples and GENCODE vM25, GRCm38, for mouse samples, and imported with tximeta^62^ for differential expression analysis with DESeq2^63^, following the recommended workflow. Genes with non-zero counts in at least two samples were retained. Genes with Padj < 0.1 and |log2FC| > 0.5 were considered differentially expressed, and shrunken log2FC values were calculated using the apeglm^64^ method.

For chromatin state enrichment analyses, the Roadmap Epigenomics^31^ 15-state core model for primary monocytes (E029) was used.

Published RNA-seq data from Dnmt3a-HET mice were obtained from NCBI GEO (accession GSE208075), comprising WT and Dnmt3a-HET differentiated macrophages under control conditions at day 8 (two samples per genotype).

### CUT&RUN

CUT&RUN was performed using the CUTANA CUT&RUN Kit (Cat # 14-1048, EpiCypher) and targeted histone mark antibodies (anti-H3K4me3 and anti-H3K27me3, EpiCypher, anti-H2AK119ub1, Cell Signaling Technology) following the manufacturer’s protocol. In brief, 0.5 million cells were harvested and nuclei were extracted. Activated Concanavalin A was incubated with nuclei for 10 min at room temperature to let the nuclei bind to the beads. For each target histone mark, 0.5 μg of H3K4me3 (Cat # 13-0041), H3K27me3 (Cat # 13-0030), or H2AK119ub1 (Cat # 8240) was added to each sample and incubated overnight at 4 °C. Isotype control (Cat # 13-0042) was used as a negative control. After overnight incubation, the beads were then washed twice with cell permeabilization buffer (Wash buffer including 0.01% digitonin), and incubated with protein AG-Micrococcal Nuclease (pAG-MNase) for 1 h at 4 °C. Excessive pAG-MNase was washed out, and then chromatin digestion was performed by adding 2 mM CaCl2. After chromatin digestion, the stop buffer (Cat # 48105, Cell Signaling Technology) and 0.5 ng *E.coli* spike-in DNA (Cat # 18-1401, EpiCypher) were added and incubated for 10min at 37 °C.

CUT&RUN libraries were prepared with the NEBNext Ultra II DNA Library Prep Kit (Cat # E7645S, NEB) and dual-indexed using Illumina adapters (NEB), following the manufacturer’s instructions. Libraries were quantified with the Qubit dsDNA HS Assay Kit (Cat # Q32851, Invitrogen^TM^, Thermo Fisher Scientific) and fragment sizes were assessed using an Agilent 2100 Bioanalyzer system. Libraries were sequenced to have at least 8 million read pairs using an Illumina NovaSeq 6000 or Illumina NovaSeq X Plus. The raw sequencing reads were first trimmed with TrimGalore v0.6.7 (https://github.com/FelixKrueger/TrimGalore) and then aligned to GRCh38 or GRCm38 using Bowtie2^65^ v2.5.1 (parameters --local --very-sensitive-local --no-unal --no-mixed --no-discordant --phred33 -I 10 -X 700). SAM files were sorted and indexed with samtools^66^ v1.15.1 to generate BAM files, and then duplicate reads were removed using GATK^67^ markduplicates (v4.2.6.1). After removing PCR duplicates and unaligned reads, bigWig files were generated from BAM files using BEDtools2^68^ v2.30 and reads were normalized by the ChIPseqSpikeInFree method^69^. Heatmap and histone mark enrichment distributions were created using deepTools^70^.

### EM-seq

Genomic DNA was isolated from hESC-derived macrophages or mBMDMs using the Quick-DNA Kit (Cat# D3020, Zymo Research). DNA were sheared to an average fragment size of ∼ 200 bp using a Covaris S2. EM-seq libraries were prepared from sheared DNA using the Enzymatic Methyl-seq kit (Cat # E7120, NEB) following the manufacturer’s instructions.

Libraries were sequenced with 150 bp paired-end reads on an Illumina platform. Quality-and adaptor-trimmed reads were aligned to the GRCh38 or GRCm38 primary genome using HISAT-3N^71^ and deduplicated using samtools^66^. CpG methylation data extracted from uniquely aligned reads using the hisat-3n-table function. Differentially methylated loci were identified using the DMLtest function in DSS^72^ with a significance threshold of p < 0.01.

### Mass spectrometry

For sample preparation, culture media was removed, and cells were washed twice with Dulbecco’s Phosphate-Buffered Saline (DPBS). Cell pellet was immediately snap-frozen in liquid nitrogen and stored at -80 °C until further processing. Histones were acid-extracted, derivatized via propionylation, and digested with trypsin, as previously described^73^. Each sample was resuspended in 300 μL of 0.1% FA/mH2O, and 2 µL was injected per run, with 3 technical replicates. Histone extraction, histone modification profiling, and mass spectrometry analysis were conducted at the Mass Spectrometry Technology Access Center at Washington University School of Medicine.

### Wound healing assay

Migration of hESC-derived macrophages was assessed using an *in vitro* scratch wound assay. Macrophages were seeded in 24-well plates and incubated for 24 h in RPMI 1640 supplemented with 10% FBS to establish a confluent monolayer. A linear scratch wound was generated across the cell monolayer using a sterile 200-µL pipette tip. Cells were gently rinsed with DPBS to remove cellular debris, and RPMI 1640 supplemented with 10% FBS was added for the duration of the assay. Images were acquired at 10× magnification at 0, 12, 24, 36, and 48 h after wounding. Wound closure was quantified as the percentage reduction in wound area relative to the wound area at 0 h using ImageJ and the Wound Healing Size Tool.^74^

For knockdown, cells were transfected with small interfering RNAs (siRNAs) using Lipofectamine RNAiMAX reagent (Cat # 13778, Invitrogen, Thermo Fisher Scientific) according to the manufacturer’s instructions. For EZH2 and RNF2 knockdown, cells were treated with 10 nM siRNAs, whereas 20 nM siRNAs were used for CXCL5 and LIF knockdown. A scrambled siRNA was used as a negative control (Cat # D-001810-10-05), and siRNAs targeting EZH2 (Cat # L-004218-00-0005), RNF2 (Cat # L-006556-00-0005), CXCL5 (Cat # L2-007879-01-0005) or LIF (Cat # L2-011720-01-0005) were purchased from Dharmacon (Horizon Discovery).

### RT-qPCR

Total RNA was isolated using Quick-RNA kit (Cat # R1051, Zymo Research). Total RNA (500ng) was reverse transcribed at 25°C for 10 min, at 37°C for 2 h, followed by 85°C for 5 min using High-Capacity cDNA Reverse Transcription Kit (Cat # 4368814, Applied Biosystems, Waltham, MA, USA). RT-qPCR reactions were prepared with PowerUp SYBR Green Master Mix (Cat # A25742, Thermo Fisher Scientific), and amplification was performed on CFX384 Touch Real-Time PCR Detection System (Bio-Rad Laboratories) or QuantStudio Real-Time PCR Systems (Applied Biosystems). Relative gene expression levels were calculated using the 2^-ΔCt^ method, with normalization to GAPDH.

### Mice

Animal experiments were performed according to a protocol approved by the Institutional Animal Care and Use Committee (IACUC) of the University of California, Irvine. Dnmt3a-R878H mice were obtained from The Jackson Laboratory (strain #032289). Hematopoietic-specific expression of the heterozygous Dnmt3a^R878H^ mutant allele was induced by crossing *Dnmt3a*^fl-R878H/+^ to *Vav-Cre* mice. Both male and female mice were used in the experiments. Genotypes were confirmed by PCR using primers 5′-CTCCTTGGATTTGAGGAGGA-3′ and 5′-TGCACATGAGAACTGGATGG-3′. The expected amplicon sizes were 282 bp for the Dnmt3a-R878H allele and 178 bp for the wild-type allele.

Bone marrow cells were isolated from the femurs and tibias of mice and cultured on untreated tissue culture plates in macrophage differentiation medium (RPMI 1640 with 10% FBS, 1% penicillin/streptomycin, and 25 ng/ mL recombinant murine M-CSF [Cat # 416-ML, R&D Systems]) for 7 days.

### Competitive bone marrow transplant

A total of 400,000 whole bone marrow cells from wild-type B6.CD45.2 or Dnmt3a-R878H mice were competitively transplanted with 1.6 × 10^6^ whole bone marrow cells from B6.CD45.1 mice at a 1:4 ratio. Cell mixtures were injected retro-orbitally into lethally irradiated B6.CD45.1/2 recipient mice. Recipient mice received 800 cGy total-body irradiation using an X-ray irradiator 16 h prior to transplantation. Recipient mice were allowed to recover for 5 weeks following transplantation. Peripheral blood (10 µL) was collected from the saphenous vein into 10 µL EDTA and red blood cells were lysed using 1 mL 1× ACK lysis buffer. Cells were washed with 1× PBS containing 2% FBS and stained with FITC-conjugated anti-mouse CD45.1 (Cat #110706, BioLegend) and PE/Cyanine7-conjugated anti-mouse CD45.2 (Cat #109830, BioLegend). Data were acquired using a NovoCyte 3000 flow cytometer (Agilent) and analyzed using FlowJo software (v10.10.0).

### Muscle injury

Prior to muscle injury, mice were anesthetized with isoflurane in oxygen. Acute muscle injury was induced by intramuscular injection of barium chloride (1.2% w/v in sterile saline). A volume of 2 µL/g body weight was injected into the quadriceps and 1 µL/g body weight was injected into tibialis anterior muscles. The contralateral quadriceps and tibialis anterior muscles served as uninjured controls. Following injury, mice were euthanized at the indicated time points, and muscles were harvested for flow cytometric analysis. Tibialis anterior muscles were harvested 7 days after injury for histological and immunofluorescence analysis.

### Muscle single cell isolation

Single-cell suspensions from mouse hindlimb muscles were generated as previously described.^75^ Briefly, mice were euthanized by carbon dioxide inhalation using a gradual-fill method in accordance with American Veterinary Medical Association guidelines and perfused with 1× PBS. Injured and uninjured quadriceps muscles were excised and mechanically and enzymatically dissociated. Cell suspensions were filtered sequentially through 70- and 40-µm cell strainers and used for flow cytometric analysis. Isolated cells were stained with Zombie NIR Fixable Viability Dye (Cat. #423105, BioLegend) for 15 min. Fc receptors were blocked by incubation with TruStain FcX anti-mouse CD16/32 antibody (BioLegend) for 15 min. Cells were stained with antibodies against CD11b (PerCP-Cy5.5, BioLegend, Cat #101228), F4/80 (PE-Cy7, BioLegend, Cat #123114), CD45.1 (APC, BioLegend, Cat #109808), and CD45.2 (PE, BioLegend, Cat #110714).

### Histological and immunofluorescence analysis of injured muscle

Tibialis anterior muscles were excised, embedded and frozen in liquid nitrogen-cooled isopentane, and stored at −80°C. Cryosections (10 µm) were prepared for immunofluorescence analysis.

For immunofluorescence staining, cryosections were fixed with 2% PFA for 5 min at room temperature and blocked with 0.2% gelatin for 30 min at room temperature. Sections were treated with 3% H2O2 and subsequently incubated with an Avidin/Biotin Blocking Kit (Cat. #SP-2001, Vector Laboratories). Following washes with PBS, sections were incubated with a biotinylated mouse antibody against embryonic myosin heavy chain (eMyHC; Cat. #sc-53091, Santa Cruz Biotechnology) and an antibody against laminin (Cat. #L9393, Sigma-Aldrich) for 3 h at room temperature. Following PBS washes, laminin was detected using donkey anti-rabbit IgG Alexa Fluor 594 secondary antibody (Cat. #A-21207, Thermo Fisher Scientific). For detection of eMyHC, sections were incubated with peroxidase-conjugated streptavidin (Cat. #016-030-084, Jackson ImmunoResearch Laboratories) for 1 h at room temperature. After washing with PBS, the eMyHC signal was amplified and visualized using iFluor 647 styramide (Cat. #45045, AAT Bioquest) for 7.5 min at room temperature using the Power Styramide Signal Amplification (PSA) system according to the manufacturer’s instructions. Sections were subsequently washed with PBS and mounted with VECTASHIELD Antifade Mounting Medium with DAPI (Cat. #H-1200-10, Vector Laboratories). Fluorescence images were acquired using a Keyence BZ-X800 microscope.

The eMyHC-positive area and total tissue area were measured using ImageJ. Images were calibrated using the scale bar before measurement. The total tissue area was manually outlined along the outer tissue boundary, and non-tissue areas resulting from sectioning artifacts were excluded to obtain the tissue area used for quantification. The eMyHC-positive area was quantified from the eMyHC channel using an intensity threshold to capture eMyHC staining while excluding background signal. The percentage of eMyHC-positive area was calculated by dividing the eMyHC-positive area by the tissue area after exclusion of sectioning-related non-tissue areas. Measurements from a single section per mouse were used to generate one value per animal, and each mouse was treated as an independent biological replicate for statistical analysis.

### Genomic and transcriptomic data availability

The RNA-seq, scRNA-seq, ATAC-seq, EM-seq and CUT&RUN data have been submitted to GEO. RNA-seq data at GEO under accession GSE289657. scRNA-seq data is available under accession GSE291728. ATAC-seq data at GEO under accession GSE289651. EM-seq data at GEO under accession GSE289660. CUT&RUN data at GEO under accession GSE289653.

## Acknowledgments

The authors thank UCI Genomics Research and Technology Hub for technical support and helpful discussions. Mass spectrometry analyses were performed by the Mass Spectrometry Technology Access Center at the McDonnell Genome Institute (MTAC@MGI) at Washington University School of Medicine. This work was funded by National Institutes of Health (NIH) R01HL153974 to M.B., B.R.R., and I.M., R21AI164367 to M.B. and I.M., P30AR070549 and CCHMC ARC Award #53632 to M.T.W., and R01AI173314 to M.B. and E.R.M.

## Contributions

Y.L., S.A.V., A.G.F., and M.B. designed experiments. Y.L., P.K.F., Y.T., J.-Y.L., X.H., C.B.,

M.E.S.W., H.-J.Y., J.L., Z.S.P., and D.J.L. performed the experiments. Y.L., P.K.F., J.P.N., X.H.,

M.E.S.W., A.N.R., A.P.P., X.C., H.Y, and M.B. analyzed data. M.T.W., I.M., B.R.R., E.R.M., E.P.,

S.A.V., A.G.F., and M.B. provided supervision. Y.L. and M.B. wrote the manuscript. All authors edited the manuscript.

---

**Extended Data Fig. 1.**
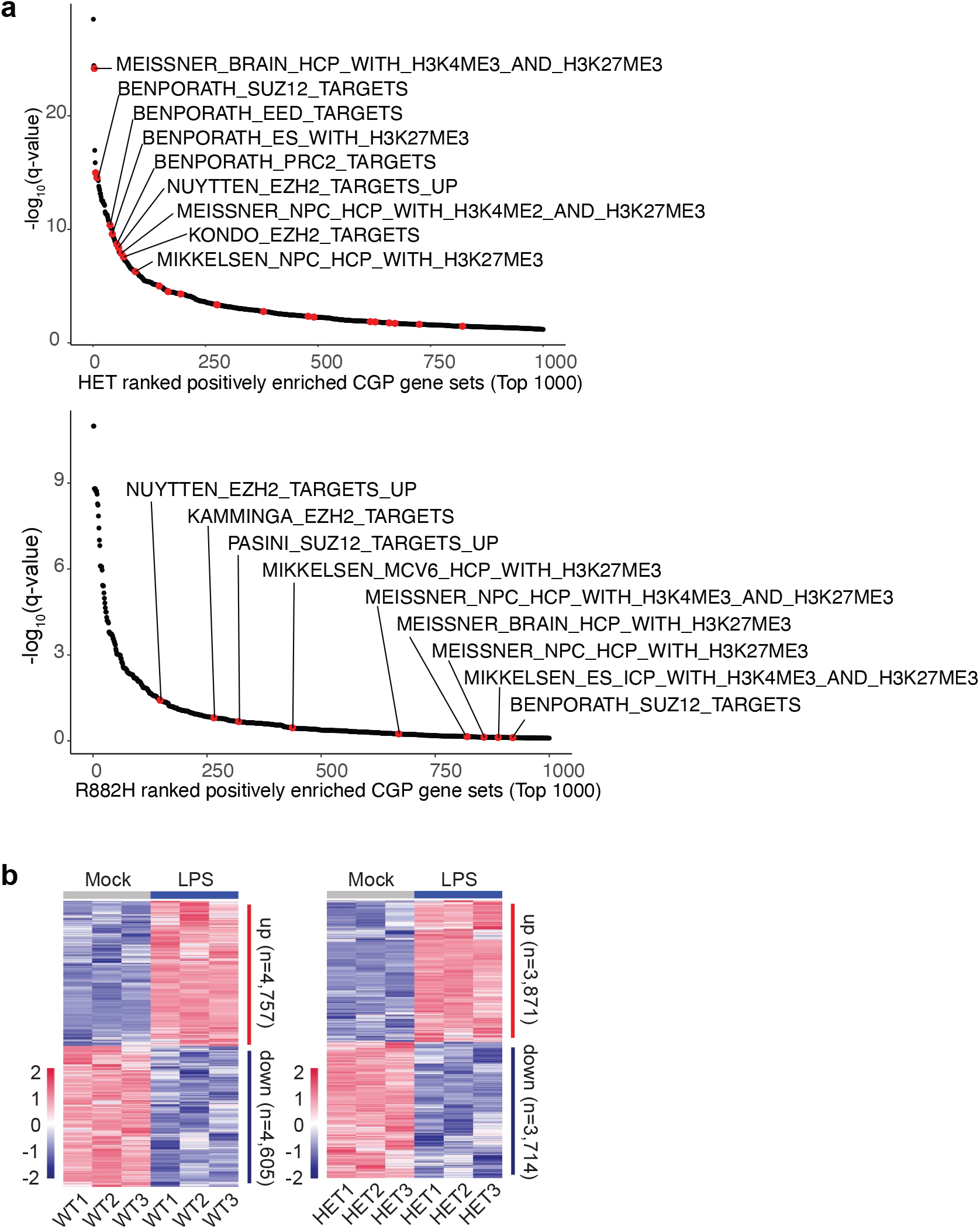
Bulk RNA-seq confirms PRC2 target gene set enrichment and reveals extensive LPS-induced transcriptional reprogramming in *DNMT3A*-mutant macrophages. **a**, Gene set enrichment analysis (GSEA) of Chemical and Genetic Perturbation (CGP) gene sets from MSigDB for bulk RNA-seq data comparing HET versus WT and R882H versus WT macrophages under steady-state conditions. PRC2-related pathways (highlighted in red) rank among the most significantly enriched recapitulating the enrichment observed in the scRNA-seq macrophage subset (Fig. 1g). Each dot represents a gene set ranked by adjusted P value (Padj) of enrichment. Top 1,000 ranked pathways are shown for clarity. **b**, Heatmap of LPS-response DEGs (LPS versus mock; Padj < 0.1 and |log2FC| > 0.5) in WT (left) and *DNMT3A*-HET macrophages (right), analyzed separately within each genotype. Rows represent DEGs and columns represent individual clones; colors indicate row-scaled expression (z-score; red, higher expression; blue, lower expression).

**Extended Data Fig. 2.**
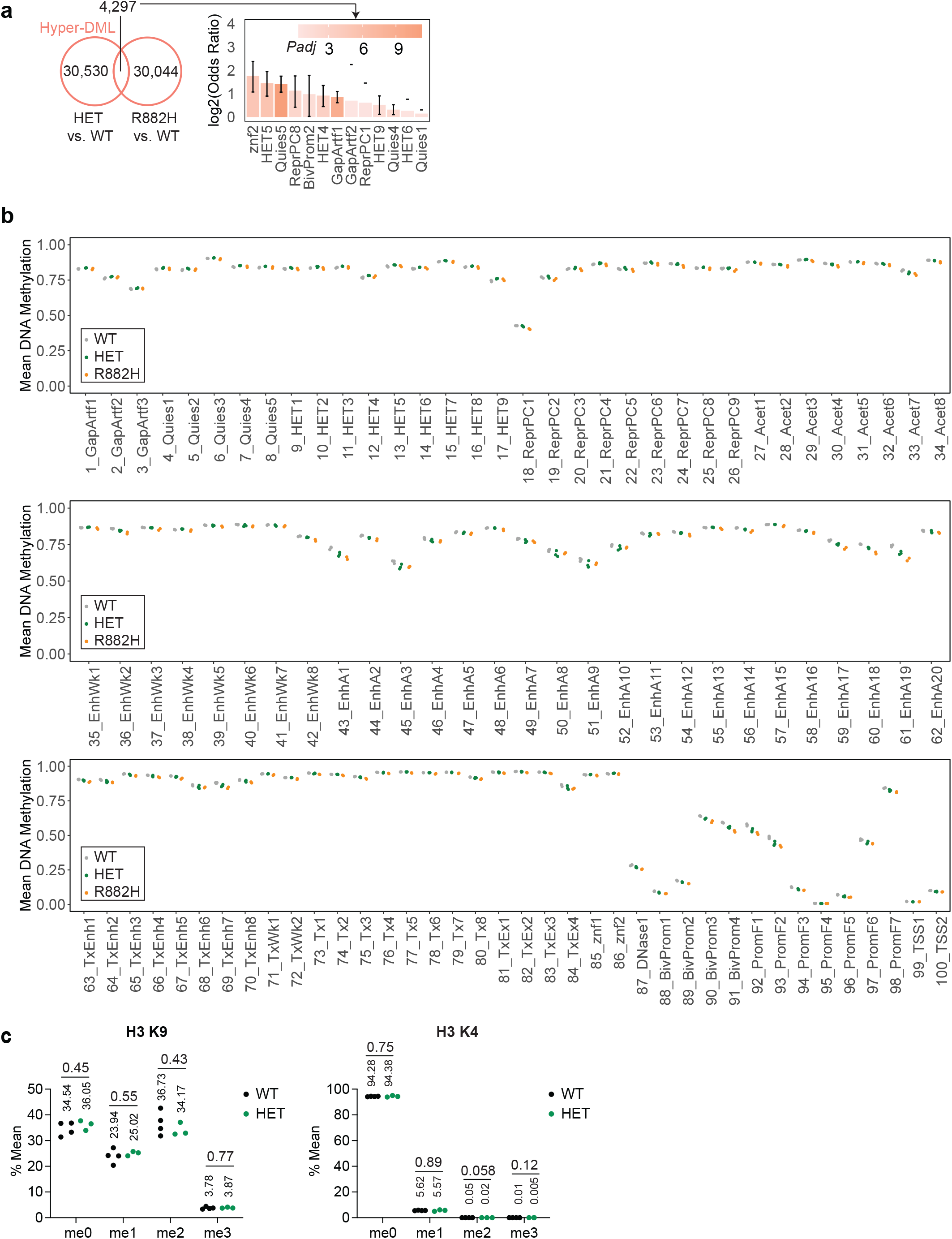
Limited enrichment of hypermethylated loci and preserved H3K9/H3K4 methylation in *DNMT3A*-mutant macrophages. **a,** Left: number of overlapping hypermethylated differentially methylated loci (hyper-DML; *DSS* DMLtest P < 0.01) between HET and R882H macrophages. Right: log2 odds ratios showing enrichment of hyper-DML shared between HET and R882H macrophages across a tissue-independent full-stack chromatin state annotation^34^. **b,** Mean methylation across CpGs within each of the 100 tissue-independent full-stack chromatin states. Each dot represents an independent clone, colored by genotype (WT, HET, R882H). **c,** Mass spectrometry quantification of H3K9 and H3K4 methylation states (me0, me1, me2, and me3) in WT and DNMT3A-HET macrophages; R882H clones were not analyzed. Values are expressed as a percentage of total H3K9- or H3K4-containing peptide. Each dot represents an independent clone. No significant genotype-dependent differences were detected. P values were calculated by Student’s t-test.

**Extended Data Fig. 3.**
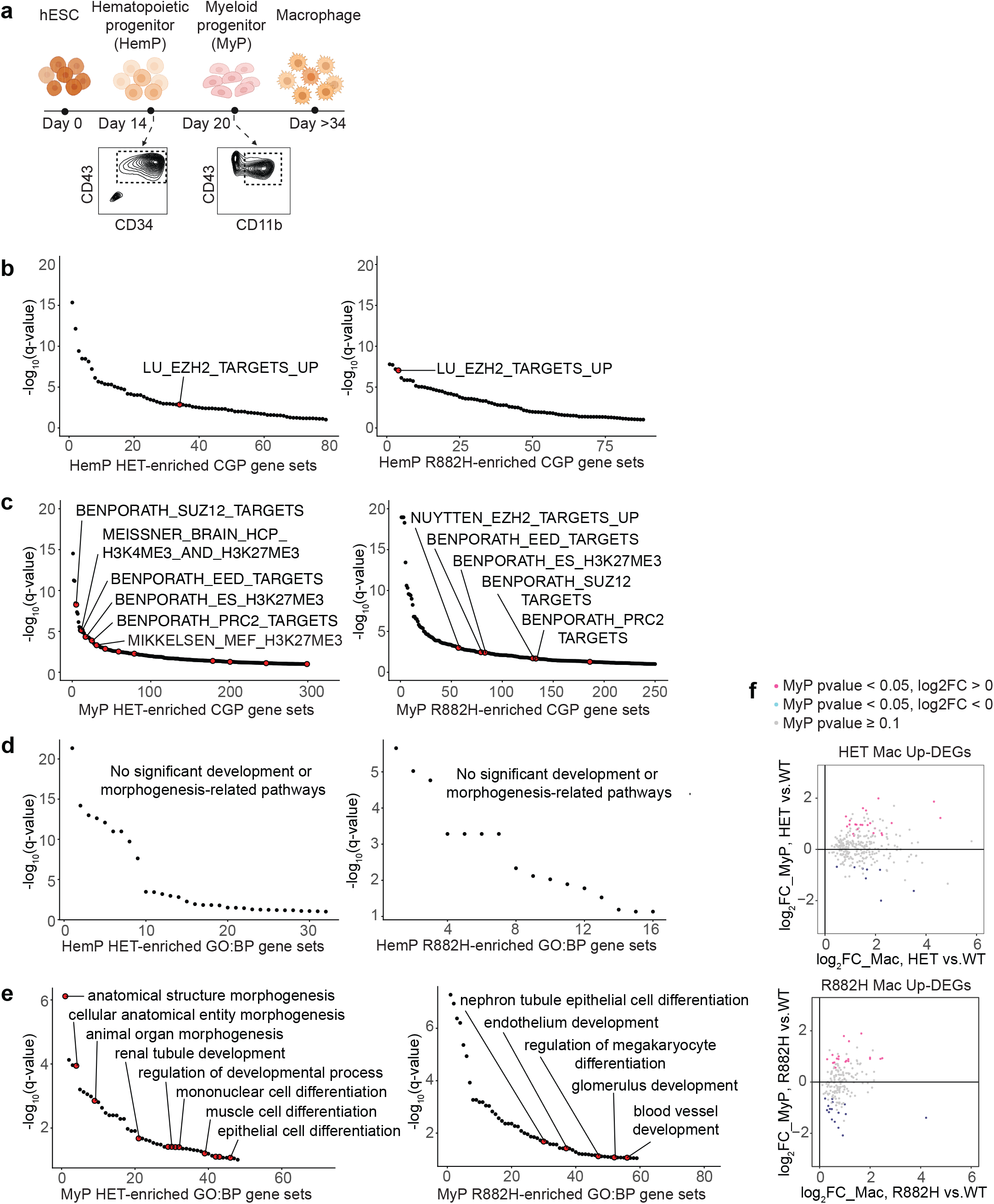
Polycomb target gene derepression emerges at the myeloid progenitor stage. **a,** Schematic of the experimental design for bulk RNA-seq of progenitor populations sorted along the *in vitro* hESC-to-macrophage differentiation trajectory: early hematopoietic progenitors (HP; CD34⁺CD43⁺, differentiation day 14) and committed myeloid progenitors (MyP; CD34⁻CD43⁺CD11b⁺, differentiation day 20), from WT, DNMT3A-HET (HET), and DNMT3A-R882H (R882H) lines. **b,** GSEA of Chemical and Genetic Perturbation (CGP) gene sets from MSigDB in hematopoietic progenitors, HET versus WT and R882H versus WT. Each dot represents a gene set ranked by Padj of enrichment; gene sets with Padj < 0.1 are shown. Only one PRC2-related gene set was found significantly enriched in both HET and R882H hematopoietic progenitors. **c,** As in b, for myeloid progenitors. Numerous PRC2-related gene sets (highlighted in red) rank among the most significantly enriched. **d,** GSEA of Gene Ontology Biological Process (GO:BP) gene sets from MSigDB in hematopoietic progenitors, HET versus WT and R882H versus WT. Each dot represents a gene set ranked by Padj of enrichment; gene sets with Padj < 0.1 are shown. No significant development- or morphogenesis-related terms were detected. **e,** As in d, for myeloid progenitors. Development- and morphogenesis-related terms (highlighted in red) rank among the most significantly enriched. **f,** Comparison of log2FC in macrophages (HET or R882H versus WT) and log2FC in myeloid progenitors (HET or R882H versus WT) for Cluster_1 (H3K27me3-high) upregulated DEGs identified in HET or R882H macrophages. Dots are colored by LPS responsiveness: magenta, LPS-induced (P < 0.05 and log2FC > 0); cyan, LPS-repressed (P < 0.05 and log2FC < 0); gray, all remaining genes.

**Extended Data Fig. 4.**
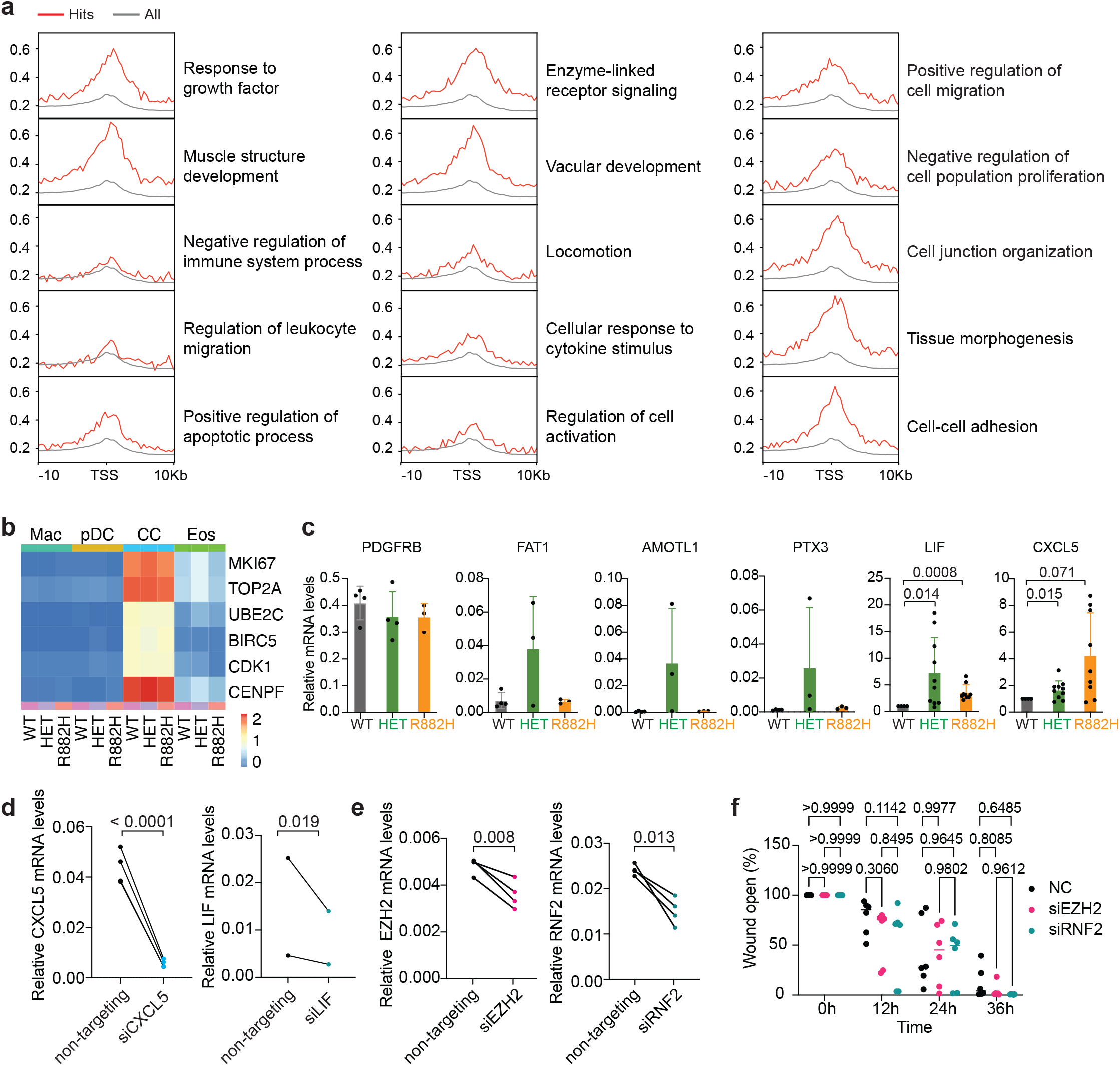
Pathway-specific H3K27me3 enrichment and competitive expansion of *Dnmt3a*-mutant hematopoietic cells. **a,** H3K27me3 enrichment analysis for Gene Ontology (GO) terms jointly upregulated in human and murine macrophages carrying DNMT3A/Dnmt3a mutations (terms as in Fig. 5a). “Hits” denotes DEGs within each GO term and “All” denotes all expressed genes in the corresponding dataset. **b**. Heatmap of proliferation-related gene expression (MKI67, TOP2A, UBE2C, BIRC5, CDK1, CENPF) across the macrophage (Mac), plasmacytoid dendritic cell-like (pDC), cycling cell (CC), and eosinophil-like (Eos) clusters in WT, HET, and R882H cells, from the scRNA-seq dataset shown in Fig. 1, with clusters defined as in Fig. 1c. Values are ln(1 + mean normalized expression), calculated per gene within each cluster and genotype. As expected, expression is highest in the CC cluster; within the macrophage cluster, WT, HET, and R882H cells show comparable levels. **c**. RT–qPCR validation of prioritized candidate genes in independent batches of hESC-derived macrophages of the indicated genotypes. Candidates were selected from Cluster_1 (H3K27me3-high; Fig. 2c) genes meeting all of the following criteria: Padj < 0.1, baseMean > 20, log2FC > 2 in HET or R882H macrophages, upregulated in both HET and R882H, and annotated to migration-related GO terms. For *PDGFRB, FAT1, AMOTL1*, and *PTX3*, values are expressed as *GAPDH*-normalized relative expression and each dot represents an independent clone. For CXCL5 and LIF, all clones — including WT — were normalized to the batch-matched WT mean (set to 1), and data from three independent batches of 3-4 clones per genotype are pooled; each dot represents one clone in one batch. P values were calculated by Welch’s t-test. **d,** *CXCL5* and *LIF* mRNA levels measured by RT–qPCR 48 h after siRNA transfection, normalized to *GAPDH*. Each dot represents an independent WT clone. P values were calculated by an one-tailed ratio paired t-test (siRNA versus siCtrl within each clone). **e,** *EZH2* and *RNF2* mRNA levels measured by RT–qPCR 48 h after siRNA transfection, normalized to *GAPDH*. Each dot represents an independent WT clone. *P* values were calculated by an one-tailed ratio paired t-test. **f.** Quantification of remaining wound area at 12, 24, and 36 h after scratch injury in WT macrophages transfected with siCtrl, siEZH2, or siRNF2. Neither siEZH2 nor siRNF2 significantly altered wound closure. *P* values were calculated by two-way ANOVA (siRNA condition × time) followed by Tukey’s multiple comparisons test.

**Extended Data Fig. 5.**
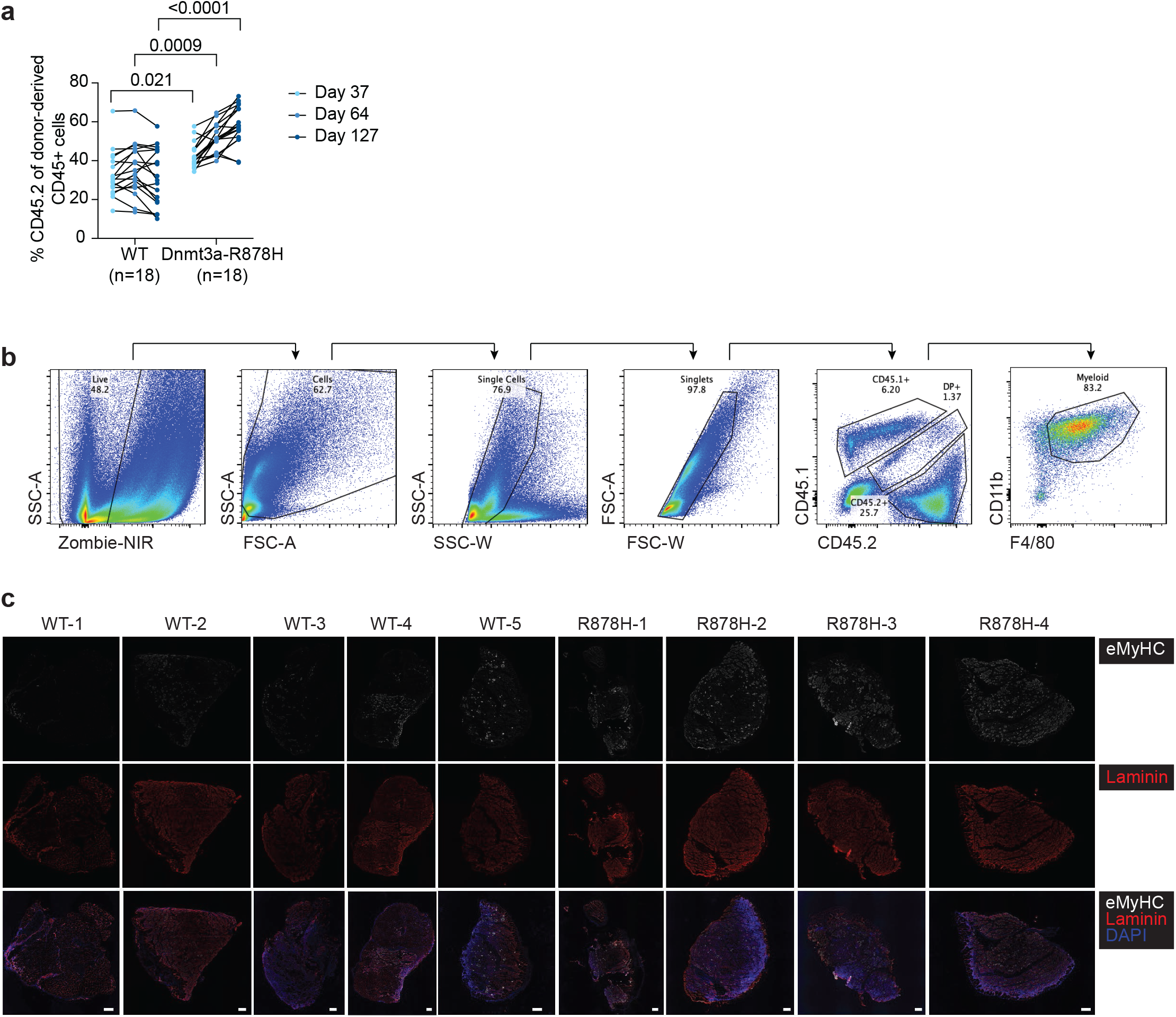
Competitive expansion of Dnmt3a-R878H hematopoietic cells and characterization of myeloid recruitment and muscle regeneration after injury. **a**, Percentage of CD45.2⁺ cells among peripheral blood leukocytes at the indicated time points after mixed bone marrow transplantation (as in Fig. 5e) in recipients of WT versus Dnmt3a-R878H test bone marrow. P values were calculated by two-way repeated-measures ANOVA (genotype × time) followed by Šídák’s multiple comparisons test at each time point. The number of mice analyzed per group is indicated. **b**, Representative sequential flow cytometry gating strategy used to identify CD45.2⁺ myeloid cells (F4/80⁺CD11b⁺) in quadriceps muscle, as quantified in Fig. 5f,g. **c**, Additional immunofluorescence images of tibialis anterior muscle sections stained for embryonic myosin heavy chain (eMyHC), laminin, and DAPI at day 7 post-injury, as in Fig. 5h. Images correspond to the mice quantified in Fig. 5i. Scale bar, 200 µm.

